# A framework for the organization of microtubules in developing neurons

**DOI:** 10.64898/2026.06.15.732274

**Authors:** Kyriacos Nicolaou, Bela M. Mulder, Lukas C. Kapitein, Florian Berger

## Abstract

The development and physiology of neurons rely on their microtubule organization, which is characterized by plus-end-out oriented microtubules in the axon and a mix of plus-end-out and plus-end-in oriented microtubules in dendrites. This orientational pattern is established early in neuronal development and is tightly linked to axon–dendrite differentiation. Even though multiple potentially relevant mechanisms have been proposed, fundamental questions remain: How does the microtubule organization in neurons emerge, and how does a neuron develop a single axon and multiple dendrites? Here, we address these questions at two distinct, complementary levels: at a higher level by proposing a conceptual framework, in which we classify mechanisms into three categories based on how they contribute to the microtubule organization: orientational bias, parallel amplification, and polarization; at a lower level we build a biophysical model that incorporates multiple mechanisms of microtubule dynamics in a neuron, from which, using analytical calculations and simulations, we derive insights into the emergence of microtubule organization in developing neurons. We show that geometrical effects alone can confer a bias in microtubule orientation. Parallel amplification then enhances the resulting polarity. Coupling multiple neurites to a common cell body that serves as a shared reservoir of resources allows for a polarization mechanism that ensures that the microtubule organization of one neurite becomes axonal while all others are dendritic. This framework unifies diverse molecular observations and yields experimentally testable predictions about microtubule self-organization in early neuronal development.

**Author summary:** Neurons communicate through long protrusions called neurites, which are of two types: dendrites, which receive signals, and axons, which send signals. Their development relies primarily on microtubules, which are polar filaments with two distinct ends, known as the plus and minus ends. Microtubules self-organize into functional architectures that are significantly different between axons and dendrites. In axons, all microtubules point their plus end away from the cell body, whereas in dendrites, they point either towards the cell body or have mixed orientations depending on the species. This orientation guides intracellular transport by motors and is closely linked to whether a neurite develops into an axon or a dendrite. Despite decades of research identifying individual mechanisms, the bigger picture behind the emergence of microtubule orientation in neurons remains unclear. Here, we construct a conceptual framework and a biophysical model to identify the principles underlying the emergence of microtubule orientation in developing neurons. Our conceptual framework provides a high-level perspective on how individual mechanisms influence microtubule organization in neurites. In our concrete biophysical model, we study a selection of mechanisms to gain specific, quantitative insight into the organizational process. We propose a minimal model of a neuron that exhibits neuronal polarization, giving rise to a single axon-like neurite and multiple dendrite-like ones, consistent with experimental observations. This *in silico* neuron helps to explain how neurons break symmetry during development and provides a systematic way to generate and test new hypotheses about neuronal polarity.

## Introduction

Neurons are building blocks of our nervous system and among the most highly polarized cells in our bodies. They receive signals with their dendrites and relay them through their axon. Establishing and maintaining this remarkable polarization and physiological function is highly dependent on the microtubule cytoskeleton [1–4]. Microtubules (MTs) are polar filaments with a distinct plus end and minus end. During the early development of a neuron, the cytoskeleton contributes to the formation of protrusions from a neuron’s cell body, called neurites. These neurites subsequently differentiate into axons and dendrites with distinct MT architecture. In axons, MTs are oriented with their plus ends pointing away from the soma, an orientation called plus-end-out (+OUT). In contrast, dendritic MTs either point with their plus-ends toward the soma, displaying a plus-end-in (+IN) orientation, or form a mixed array of both orientations, depending on the species [5, 6]. In addition to the different MT organization in axons and dendrites, the abundance of microtubule-associated proteins (MAPs) also differs between these compartments [7–11]. These differences in MT organization between the axon and the dendrites are likely linked to their distinct physiological processes, such as directed cargo transport, positioning of organelles, and the formation of synapses [10, 12, 13]. Despite decades of research, a central question remains: How do developing neurons self-organize their MT cytoskeleton to generate these distinct fate-determining patterns?

To answer this question, we must look at the early stages of neuronal development, when neurites begin to grow and differentiate into axons or dendrites [3, 14]. In stage 1, protrusions rich in filamentous actin emerge from the soma. In stage 2, the protrusions develop into undifferentiated neurites as their cores become progressively populated with MTs. In stage 3, one neurite outgrows the others and becomes the axon. In stage 4, the remaining neurites differentiate into dendrites. Throughout stages 2 and 3, the MT cytoskeleton undergoes intense remodeling: individual MTs stochastically grow and shrink, displaying dynamic instability, slide in and out of the neurite by motor proteins [15–18], nucleate from non-centrosomal sites [19–21], and become decorated by MAPs [11, 22, 23]. Remarkably, perturbing these processes, for example, by stabilizing MTs, can disrupt the polarization of the neuron, leading to neurons developing multiple axons [16, 24, 25], indicating that neurite fate follows from MT organization and not vice versa.

The extensive remodeling of the MT cytoskeleton during stages 2 and 3 has attracted significant attention from both experimental and theoretical communities, in search of the mechanisms underlying the emergence of neuronal polarity [1–4, 26–28]. While axon specification can be biased by extracellular cues [16, 29–32], neurons are capable of polarizing in their absence, suggesting that intrinsic mechanisms are sufficient to establish polarity [16, 33]. A prevailing view is that once one neurite begins to outgrow the others, a positive feedback loop reinforces its identity as the future axon, partly caused by the accumulation of +OUT MTs [34–36]. This uniform +OUT MT orientation could facilitate active transport of growth-promoting factors by motors toward the neurite tip, thereby enhancing neurite elongation. Thus, neuronal polarization closely fits a classical local-excitation / global-inhibition (LEGI) scheme: local positive feedback amplifies a slight random bias in one neurite, while slower, system-wide negative feedback, such as the depletion of a shared factor, suppresses the same response in the other neurites. Small GTPases of the Rho and Ras families instantiate this scheme and are important effectors in the symmetry breaking that selects a single axon among the developing neurites [37–41]. The resulting neuron has a single axon with a uniform +OUT MT array and multiple dendrites with mixed or uniform +IN MT arrays, aligning neuronal symmetry breaking with other well-studied polarity systems [42, 43]. While this scheme is compelling at an abstract level, the detailed molecular mechanisms underlying the formation of the dendrites’ and the axon’s specific MT organization remain unclear.

A wide variety of individual mechanisms have been implicated in the establishment and maintenance of MT polarity in developing neurons. Motor-driven sliding can sort MTs, with dynein removing +IN MTs from axons [17, 44], and kinesin-1 transporting +OUT MTs out of dendrites [17, 45, 46]. Non-centrosomal nucleation, often centered around Golgi outposts or induced by the augmin complex, establishes +IN MTs into dendrites and +OUT MTs into axons [19–21]. Severing enzymes, like katanin and spastin, selectively amplify correctly oriented MTs [47]. Live imaging further reveals orientation-dependent dynamics: +OUT MTs in the distal axon grow more persistently than +IN MTs [36]. While each of the proposed mechanisms captures a facet of MT organization, studying them in isolation ignores the complex interactions and feedback among them [48].

Here, we aim to understand how MTs self-organize in developing neurons using two complementary approaches. First, we construct a conceptual framework that provides a unified way to interpret the effects of individual mechanisms already suggested in the literature, and to understand how they work together to enable the emergence of the MT organization. Instead of considering a wide range of detailed mechanisms, we base our analysis on three classes: orientational bias, parallel amplification, and polarization. We argue that different molecular mechanisms can be categorized into these three classes, thus providing a systematic framework for understanding MT organization in developing neurons. Moreover, to address specific open questions about MT organization, we introduce a biophysical model that enables systematic study of selected mechanisms. First, we consider the case of a single neurite that forms a half-closed compartment. Using analytical methods and stochastic simulations, we study how different effects of our model contribute to MT organization. With this approach, we develop an intuitive understanding of different MT organizations in half-closed compartments. By coupling multiple neurites to a common cell body, we establish a model for a whole neuron. In this model, the symmetry between neurites breaks such that a single axon-like neurite emerges while the rest adopts a dendrite-like identity.

## Results

### Conceptual framework: Three classes of mechanisms shaping the microtubule organization in developing neurons

The emergence of the MT organization in developing neurons is a highly complex process, influenced by numerous mechanisms. We take a general perspective in which we classify relevant mechanisms into three distinct classes, which we refer to as: orientational bias, parallel amplification, and polarization, see Fig 1A-C. Each class captures a distinct fundamental mode of the MT-organization machinery of neurons. Specific molecular mechanisms can be categorized into these classes, thereby reducing highly complex processes to their effective influence on MT organization. Below, we define these classes and explain how specific experimental observations fall into each category.

**Fig 1.**
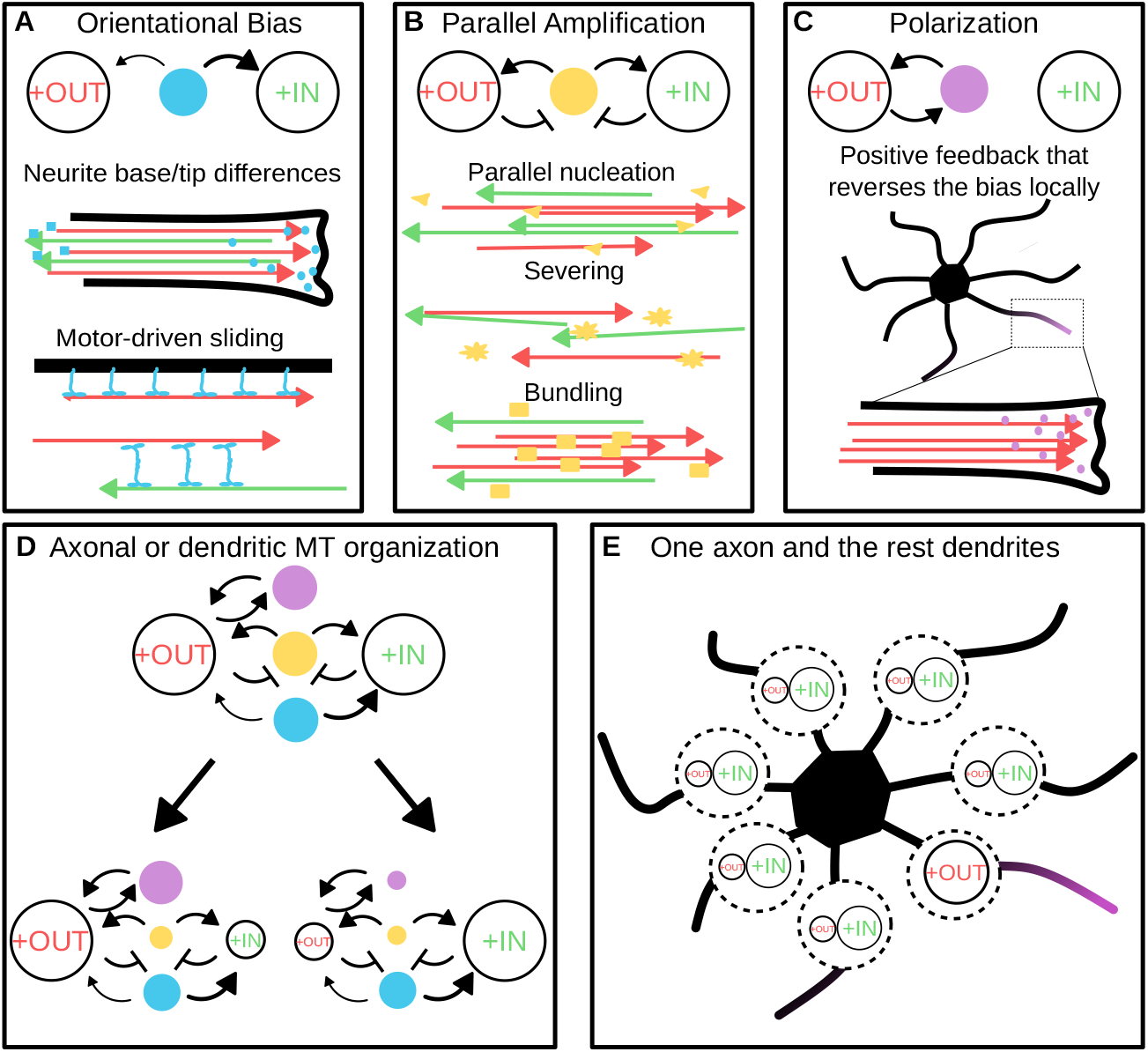
Three mechanisms underlying the emergence of the microtubule organization in developing neurons. (A) Orientational bias favors one MT orientation over the other: red arrows illustrate +OUT MTs and green arrows +IN MTs. This bias results from the neurite’s geometry, as MTs in the two possible orientations experience different biochemical and biophysical environments at both ends. Also, motor-driven sliding by motors can contribute to orientational bias. (B) Parallel amplification involves mechanisms that increase the number of MTs based on preexisting orientation through reactions such as parallel nucleation, severing, and bundling. (C) For polarization, a polarity factor (magenta) accumulates in the neurite that transiently contains more +OUT MTs, stabilizes those MTs, and closes a positive-feedback loop that can reverse the bias locally. (D) The interaction of these mechanisms determines the fate of a single neurite: an axon-like state dominated by +OUT MTs or a dendrite-like state with mixed or +IN arrays. (E) Under the right conditions, coupling a few neurites to form one neuron with a shared polarity factor ensures the correct emergence of the MT organization. The polarity factor is shared through the soma, captured by one neurite, and depletes it from its neighbors; the resulting local-excitation/global-inhibition scheme ensures that exactly one axon emerges while the remaining neurites adopt dendritic identities.

#### Orientational bias

Orientational bias accounts for mechanisms that selectively promote one orientation over the other, see Fig 1A. Such mechanisms actively discriminate one MT orientation, for example, by enhancing +OUT MTs and diminishing +IN MTs, or vice versa. We argue that these effects are primarily geometrical in origin, as +OUT MTs and +IN MTs are exposed to different environments due to the morphology of the neurite; one end of a neurite is connected to the cell body, and the other to the growth cone. These mechanisms often involve motor proteins that slide MTs in opposite directions depending on their orientation. For example, cortically anchored dynein can pull +IN MTs out of the neurite while reeling in +OUT MTs from the soma, see for example [17, 49]. Beyond motor proteins, the geometry of the neurite itself can create differential growth environments: plus ends of +OUT MTs may experience distinct regulatory factors or signaling molecules at the neurite tip, while plus ends of +IN MTs are exposed to the environment of the cell body. Thus, gradients within the neurite lead to asymmetrical rates of growth or degree of stabilization [36, 50]. Together, these directional influences establish a default orientational bias, which can be further enhanced by other mechanisms, such as parallel amplification.

#### Parallel amplification

By parallel amplification, we refer to processes that amplify the number or total mass of MTs of a particular orientation, without inherently favoring +OUT over +IN MTs, see Fig 1B. For example, MAPs such as TRIM46 [11, 23] or Tau [51] bundle parallel MTs, which often leads to collective stabilization. Severing enzymes like katanin or spastin can cut a single MT into two of the same orientation, effectively increasing the number of MTs with that orientation [52, 53]. Augmin-*γ*-TuRC complexes nucleate MTs by copying the orientation of others, leading to amplification of a dominant orientation with respect to the other [19, 20, 54, 55]. Because these amplifying mechanisms reinforce any initial bias, a neurite has a tendency to converge to uniform polarity with all +IN or all +OUT MTs. However, parallel amplification alone does not determine which orientation dominates; it merely amplifies whichever orientation is already present.

In developing neurons, where many neurites adopt dendritic identity and only one becomes the axon, it is reasonable to consider that the dendritic MT architecture is the default one, with the axonal MT architecture being the exception to the rule. Thus, the directional bias, enhanced by parallel amplification, in all neurites favors +IN over +OUT. For the emergence of an axon-like orientation with all +OUT MTs, a polarization mechanism is needed to tilt the balance in favor of a +OUT orientation.

#### Polarization

Polarization is a mechanism by which a single neurite locally overrides the default dendritic bias towards an axonal one, see Fig 1C. It should involve a positive feedback loop: a slight excess of +OUT MTs in one neurite attracts proteins that further stabilize and expand that orientation, thereby triggering a *winner-takes-all* outcome. This requirement for orientation selection could, for example, be met through the upstream involvement of motor proteins such as kinesin-1. These motors preferentially move into the neurite that has more +OUT MTs and, once there, may destabilize +IN MTs or stabilize +OUT MTs [36, 42, 56]. Such positive feedback creates a self-reinforcing cycle, in which the neurite that is best suited to become the axon retains and amplifies its +OUT MTs. In contrast, the other neurites remain susceptible to any default bias, which ultimately results in a mixed-polarity dendritic architecture. Thus, polarization ensures that only a single neurite commits to a strong +OUT orientation, allowing it to develop into the axon.

#### Microtubule organization in developing neurons: an interplay of three classes of mechanisms

In summary, this triad of classes provides a consistent conceptual framework for understanding how neurons achieve their distinct axonal and dendritic MT organization, see Fig 1D. Directional bias sets a baseline preference for either +OUT or +IN orientation, and parallel amplification magnifies this orientation. For a neuron to specify a single axon, polarization acts as a winner-takes-all mechanism, see Fig 1E.

This framework provides conceptual insight into how different mechanisms shape MT organization. To obtain quantitative predictions, we construct a biophysical model based on a selection of mechanisms whose outcomes we interpret in the context of our framework.

### Biophysical model of a single neurite

We build a biophysical model for the organization of MTs in neurons in two steps. First, we consider a single neurite in which MTs display dynamic instability, sliding, parallel nucleation, transition to a boundary state at the tip, and entry from the cell soma. The single neurite description aims to investigate how these effects differentially contribute to orientational bias and parallel amplification. In the second step, we consider a whole neuron with several neurites coupled by a cell body that acts as a reservoir for shared factors. The MTs in the individual neurites of the neuron undergo the same dynamics as introduced in the single neurite description, but modulated by a shared pool of polarity and growth factors.

#### A biophysical model of microtubule organization in a single neurite

We describe a neurite as a one-dimensional compartment of length *L* with spatial coordinate *x* ∈ [0, *L*], extending from the soma junction at *x* = 0 to the tip of the neurite at *x* = *L*, see Fig 2. MTs are represented as oriented line segments characterized by their two distinct ends, their minus end and plus end. We track their position along the neurite, using the location of their minus end, *q* ∈ [0, *L*], as the reference point. Their orientation is specified as +OUT or +IN. Throughout, we use a right-pointing arrow 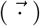 above symbols to indicate that a quantity is associated with the +OUT orientation and a left-pointing arrow 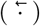 for an association with the +IN orientation. Finally, we use the length *l* ∈ ℝ^+^ of MTs to explicitly account for dynamic instability.

**Fig 2.**
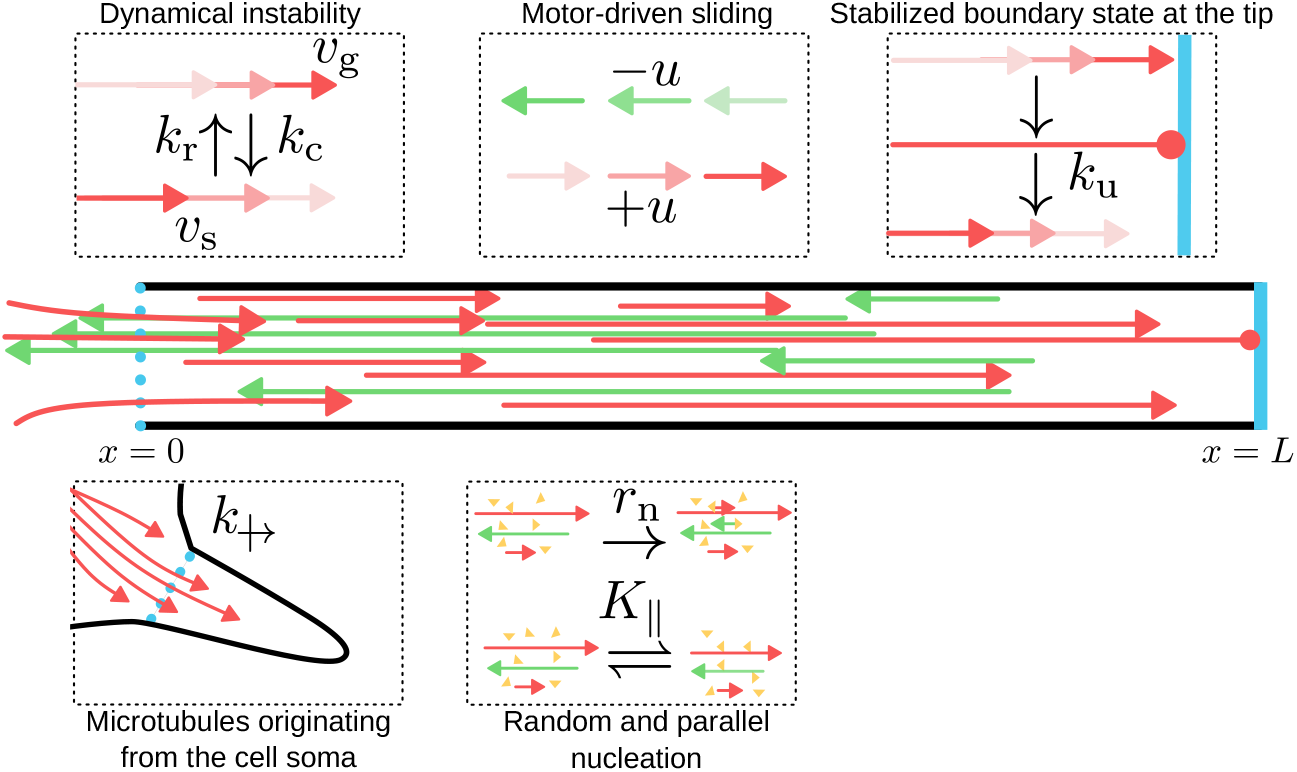
Biophysical model of MT dynamics in a single neurite. In the middle: We consider a neurite as a one-dimensional compartment populated by MTs oriented either +OUT (red arrows) or +IN (green arrows). The neurite is open at the base, at *x* = 0, to the cell soma and closed at its tip, at *x* = *L*. In the dashed boxes, we illustrate the model’s different mechanisms. In the top row, from left to right: (i) *Dynamic instability:* MTs grow with speed *v*_g_ and shrink with speed *v*_s_. They transition between those states with catastrophe rate *k*_c_ and rescue rate *k*_r_. (ii) *Motor-driven Sliding:* +OUT MTs and +IN MTs slide in opposite directions with speed *u*. (iii) *Stabilized boundary state:* When +OUT MTs reach the tip at *x* = *L* (blue line), they transition immediately into a boundary state which they exit with rate *k*_u_. In the bottom row from left to right: (i) centrosomal MTs originating from the cell soma cross the neurite junction with rate 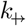 (ii) Nucleating factors (yellow triangles) can be free or bound to MTs with a binding constant *K*_∥_, and can spontaneously nucleate MTs with rate density *r*_n_.

#### Measures of microtubule organization in a neurite

We characterize the organization of a collection of MTs by the joint distribution of minus-end locations, lengths, and orientations. However, such a description is rather difficult to interpret and compare, necessitating the construction of simpler measures of organization. As we are predominantly interested in the orientation of MTs, we will quantify the organization of MTs by the polarity profile, which we define as

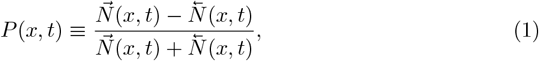

which is the normalized difference between the number 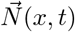 of +OUT and the number 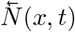 of +IN MTs crossing location *x* at time *t*. We refer to these numbers as the MT *crossing numbers*. The local polarity defined by Eq (1) attains values between 1, indicating a uniform +OUT organization, and − 1, indicating a uniform +IN organization. By spatial averaging, we define the mean polarity as

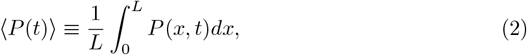

which quantifies the polarity along the entire neurite with a single number, making it particularly convenient for comparisons between multiple neurites.

#### Densities of microtubule subpopulations

We distinguish two subpopulations of MTs: (i) The *non-centrosomal* MTs that are nucleated within the neurite, labeled by n, which we describe by the orientation-resolved joint densities 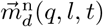 for the +OUT orientation and 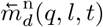 for the +IN orientation as functions of their minus-end locations *q* ∈ [0, *L*] and their length *l*. Here, the subscript *d* denotes the *dynamical state* of their plus end, which is further described below. (ii) The *centrosomal* MTs that are nucleated within the soma, labeled by c. As these MTs enter the neurite at *x* = 0, they can only be +OUT and are described by the densities 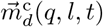, with *q* ∈ (−∞, 0], where for convenience we disregard the finite size of the soma. The dynamical state *d* of MTs can be *growing, d* = g or *shrinking, d* = s. The plus ends of +OUT MTs can, in addition, reach the tip of the neurite at *x* = *L*, in which case they may enter a *bound* state *d* = b. The dynamics of these densities will be described in the following section.

In discussing the total number of MTs of a given orientation, it is convenient to introduce the following convention. When either the superscript–indicating subpopulations–or the subscript–indicating dynamical states–is omitted, summation over the relevant categories is implied. Thus, for the +OUT MTs we have

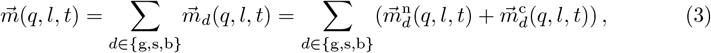

and for the +IN MTs

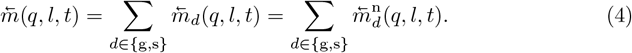

To determine the polarity measures, we need to derive the MT crossing numbers 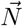 and 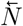 from the *m*-densities. A +OUT MT crosses a location *x* in the neurite, if its minus end, located at *q*, and its plus end, located at *q* + *l*, bracket *x*, i.e., *q < x < q* + *l*. For a given minus-end location *q*, we need to integrate 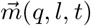 over all lengths *l* from *x* − *q* to *L* − *q*, with the upper bound being the maximum length a +OUT MT can have, as the neurite is closed at the tip. Then, we integrate over all potentially contributing MT minus-end positions *q*, where to account for the possibility that non-centrosomal MTs slide out of the neurite, and to include centrosomal MTs, we extend the integration interval with the interval −∞ *< q <* 0. Thus, 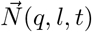 follows from 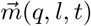 by the following integral

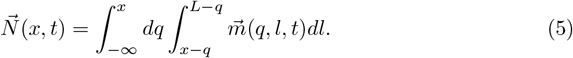

Similarly, we derive the crossing number for +IN MTs by again considering that the minus end, located at *q*, and the plus end, located at *q* − *l*, must bracket *x*, i.e., *q* − *l < x < q*. The *l*-integration is then taken over the interval *q* − *x < l <* ∞, which again assumes that MTs can grow indefinitely far into the soma, and the *q* integration runs from *x* to *L*. Using these bounds, we obtain 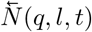 from 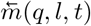 as

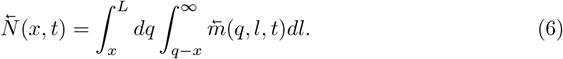

#### Dynamics of MT subpopulations

We now introduce our model for MT organization within a single neurite by specifying the dynamics for the different MT subpopulations.

#### Non-centrosomal MTs

For the length dynamics of MTs in the growing or shrinking state, we use the classic Dogterom–Leibler two-state model for dynamic instability [57]. Thus, when a MT’s plus end is in the growing state *d* = g, it gains length with speed *v*_g_, and in the shrinking state *d* = s, it loses length with speed *v*_s_. The transition from the growing state to the shrinking state occurs with the catastrophe rate *k*_c_ and the reverse transition with the rescue rate *k*_r_.

Once the plus end of a +OUT MT reaches the boundary at the tip of the neurite, its dynamical state transitions into a boundary state *d* = b. In this boundary state, the MT is stabilized, and its plus end is stalled at the boundary. Bound MTs leave the boundary state with rate *k*_u_ into the shrinking state.

Finally, non-centrosomal MTs may be subject to an effective sliding mechanism. The sliding speed is independent of orientation and is denoted by *u*. However, the direction of sliding does depend on orientation: for positive *u*, +OUT MTs move away from the soma while +IN MTs move towards it, and for negative *u* the other way around.

The time evolution of non-tip-bound, non-centrosomal +OUT MTs is governed by

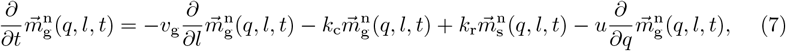

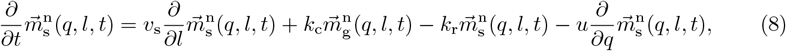

and correspondingly the +IN ones by

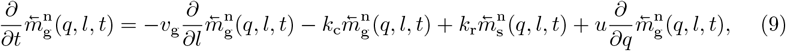

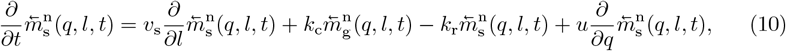

in which the first three terms on the right-hand side describe the length change and the transitions from and to the shrinking and growth states. The last term is an advection term that accounts for sliding.

The +OUT MTs reach the neurite tip with their plus ends when their length equals *l* = *L* − *q*. There, they transition to the boundary state *d* = b. The dynamics of the boundary population is described by

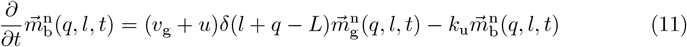

with the first term on the right-hand side being the influx of growing MTs into the boundary state and the second term the outflux at rate *k*_u_. Here, *δ*(*x*) is the Dirac delta function. Additionally, the transition from the boundary state into the shrinking state implies the boundary condition

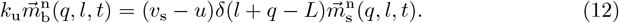

There are a few additional boundary effects: (i) Our model neurite has an open boundary at the soma–neurite junction for *x* = 0. Here, +IN MTs can freely grow and slide into the cell soma. However, if their minus ends slide beyond *x* = 0, they leave the system. Similarly, the minus end of +OUT MTs can cross *x* = 0 for *u* < 0, and they leave the system if the plus end also leaves the neurite domain *q* + *l* < 0. (ii) In case *u <* 0, the minus end of a sliding +IN MT can meet the tip boundary when *q* = *L*, we then simply consider that the sliding halts.

#### Dynamics of centrosomal MTs

Centrosomal MTs enter the neurite at *x* = 0. We assume that the total rate of growing centrosomal MTs that cross the neurite base is constant and given by 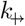. Therefore, we introduce an effective nucleation rate at the neurite base as

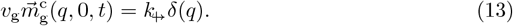

Thus, we count their length only within the neurite and effectively assign their minus-end position at *q* = 0. Note that this approach implicitly assumes that the centrosomal MTs that lie entirely in the soma are always in equilibrium, so that 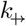 is independent of time. Moreover, it implies that we are agnostic about the position *q* < 0 in the soma where these MTs are nucleated. This requires an appropriate adjustment in the application of Eq. (5) to the calculation of the +OUT crossing number 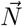.

Otherwise, centrosomal microtubules are always +OUT and undergo the same dynamical effects as non-centrosomal ones, with the exception that their minus ends are fixed and therefore cannot slide. Thus, this subpopulation is governed by the dynamical equations

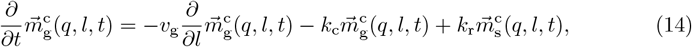

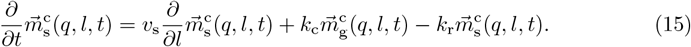

Again, similar to non-centrosomal MTs, once the plus end of a centrosomal MT reaches the tip of the neurite, it transitions into the boundary state whose population density then follows

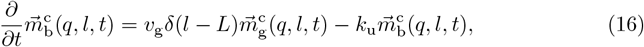

with the transition from the boundary state to the shrinking state implying the condition

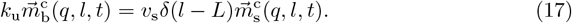

#### Nucleation of new non-centrosmal MTs

Finally, we consider the mechanism by which non-centrosomal MTs are nucleated within the neurite. To that end, we assume a finite number of *nucleators* present in the neurite. These nucleators can be bound to MTs present in the neurite or can freely diffuse within its volume. Assuming that both diffusion and the binding/unbinding kinetics of nucleators are fast compared to the timescale of the MT dynamics, we can describe the balance between bound and unbound nucleators as being in quasi-equilibrium. Free nucleators nucleate MTs with equally likely orientations, while bound nucleators nucleate MTs with the orientation of the MT they are bound to. This nucleation mechanism will act as the parallel amplifier of our system.

In Materials and Methods Parallel nucleation kinetics, we derive from first principles the nucleation rate densities for the two orientations using the assumptions above. The result is

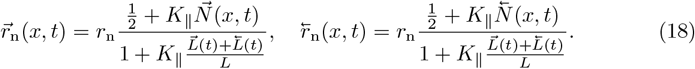

Here *r*_n_ is the constant total nucleation rate density, i.e, *r*_n_*L* is the total nucleation rate, *K*_∥_ the equilibrium binding constant of the nucleators, and 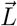 and 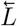 the total length of +OUT and +IN MTs respectively. For vanishing parallel nucleation, *K*_∥_ = 0, the nucleation rate densities simplify to 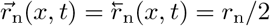, implying uniform nucleation across the neurite and equally probable in both orientations.

Using the nucleation rate densities from Eq (18), we write the nucleation conditions for the densities of growing non-centrosomal MTs for *l* = 0, as

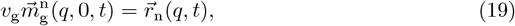

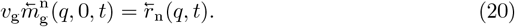

#### Approaches

To summarize, we describe the MT dynamics by the *m*-densities, from which we then derive the crossing numbers 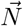 and 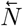. These numbers define the polarity profile and the mean polarity. This approach allows us to study how different mechanisms affect the MT organization in a single neurite. In Derivation of analytical solutions and Derivation of semi-analytical solutions for non-zero *K*_∥_, we derive expressions for the *m*-densities, crossing numbers, and polarity, for special cases of the model; for the most general case, we resort to direct simulations of the stochastic processes underlying the model, see Event-driven simulations. In Numerical values for model parameters, we make an analysis to provide reasonable numerical values and ranges for the parameters of our model so that the emergent MT organization is consistent with experimental observations from literature.

#### Spatial boundaries induce orientational bias of MT organization

In Derivation of analytical solutions, we solve the equations of the model at steady state for the special case in which nucleation factors do not bind to MTs and MTs do not slide, *K*_∥_ = 0 and *u* = 0. We obtain closed-form expressions for the MT densities, the MT crossing numbers, and the polarity profile. The closed-form expressions for the MT crossing numbers for both orientations read

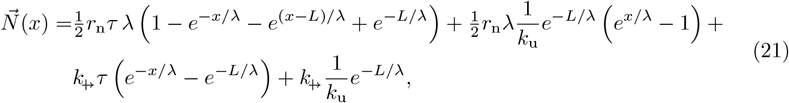

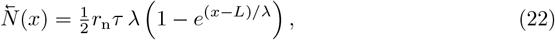

in which

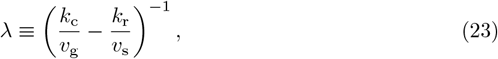

and

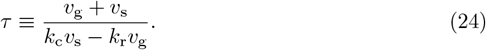

We identify *λ* as the mean length of MTs undergoing dynamic instability in an unbounded space, and *τ* as their lifetime, previously introduced in [57]. In the following, we refer to these quantities as the mean length and lifetime, although within the spatial constraints of our model, they do not represent the actual mean length and lifetimes of the MTs. Throughout this paper we choose a fixed set of numerical values for the dynamical-instability parameters, presented in Table 1, which correspond to *λ* = 7.9 µm, and *τ* = 109 s calculated from Eq (23) and Eq (24) respectively, see also Numerical values for model parameters.

**Table 1.** Numerical values for parameters describing the dynamic instability of MTs used throughout this work, and the corresponding mean length *λ* calculated from Eq (23) and lifetime *τ* from Eq (24).

| Parameter | Description | Value |
| --- | --- | --- |
| $v_g$ | growth speed | $0.10 \mu\text{m s}^{-1}$ |
| $v_s$ | shrink speed | $0.27 \mu\text{m s}^{-1}$ |
| $k_c$ | catastrophe rate | $0.12 \text{s}^{-1}$ |
| $k_r$ | rescue rate | $0.29 \text{s}^{-1}$ |
| $\lambda$ | mean length | $7.9 \mu\text{m}$ |
| $\tau$ | lifetime | $109 \text{s}$ |

To investigate the implications of the mechanisms in our model, we explore four scenarios, each realized with a different set of numerical values for the model parameters. The numerical values are summarized in Table 2 and the corresponding polarity profiles are plotted in Fig 3.

**Table 2.** Numerical values for parameters used to explore four different scenarios shown in Fig 3.

| Parameter | Scenario 1 | Scenario 2 | Scenario 3 | Scenario 4 |
| --- | --- | --- | --- | --- |
| $r_n (\mu\text{m}^{-1} \text{s}^{-1})$ | 0.05 | 0.05 | 0.05 | 0.05 |
| $L (\mu\text{m})$ | 1.0, 10, 100 | 10 | 10 | 10 |
| $k_{\text{+}} (\text{s}^{-1})$ | 0 | 0.0, 0.2, 0.4 | 0.04 | 0.04 |
| $k_{\text{u}} (\text{s}^{-1})$ | ∞ | ∞ | 0.001, 0.01, 0.1 | 0.01 |
| $u (\mu\text{m} \text{s}^{-1})$ | 0 | 0 | 0 | -0.03, 0.0, 0.03 |

**Fig 3.**
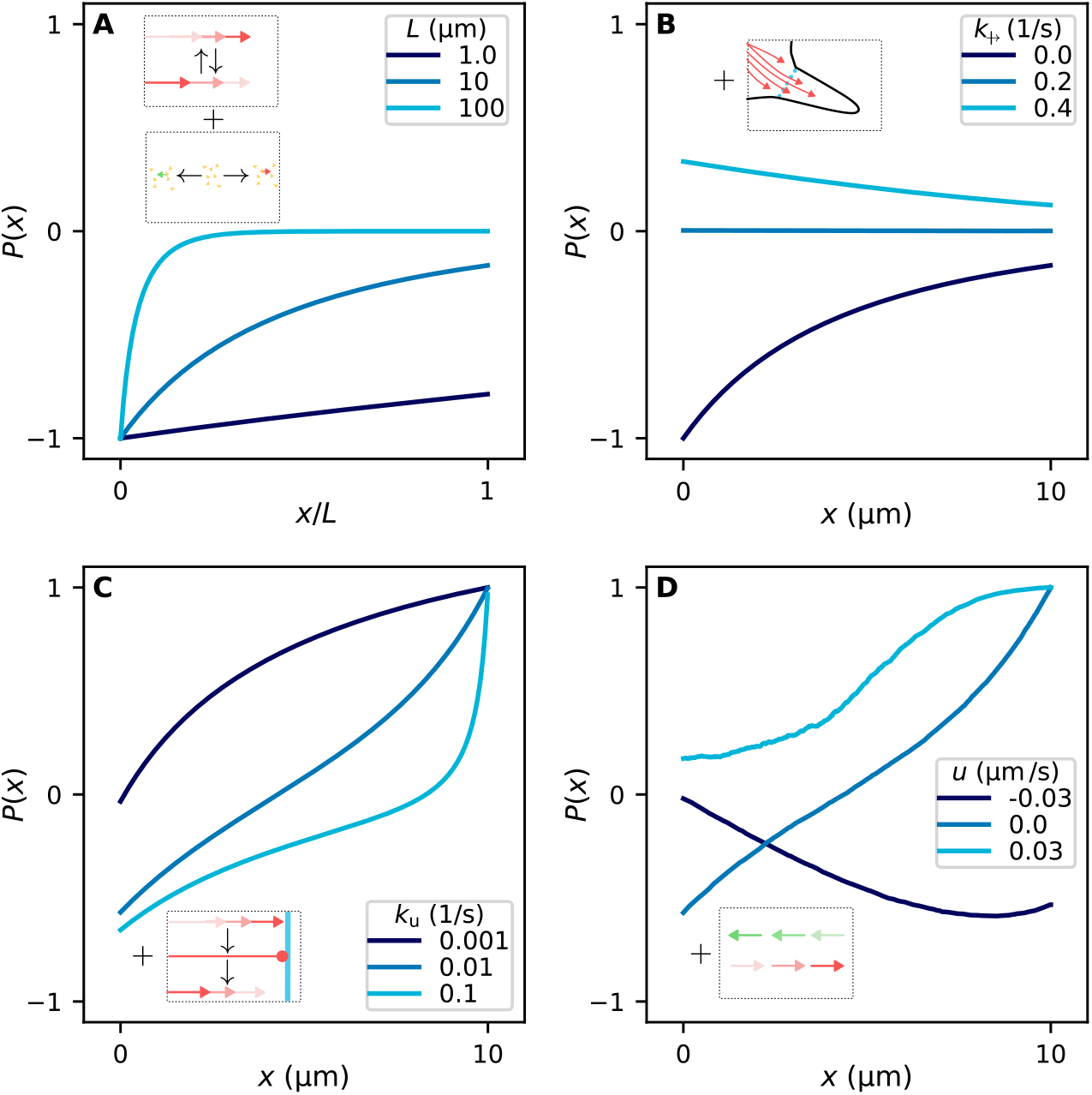
The neurite geometry induces an orientational bias by spatially asymmetric mechanisms. Polarity profiles *P*(*x*), where *x* is the coordinate along the neurite. Each panel corresponds to a scenario in which we include an additional mechanism from A to D, illustrated in the insets. (A) the baseline model in which MTs nucleate uniformly and equally likely with both orientations and undergo dynamic instability, with growing MTs shrinking upon contact with the neurite’s tip. We show the polarity profile from Eq (25) for different neurite lengths *L*. (B) We add the crossing of centrosomal MTs from the soma to the mechanisms of A. The polarity profile from Eq (26) attains positive and negative values depending on the rate 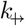 of centrosomal MTs entering the neurite. (C) Additionally, we consider that MTs change into a boundary state at the tip. They leave this boundary state with rate *k*_u_. The polarity profile *P*(*x*) from Eq (72) can change sign along the neurite depending on *k*_u_. (D) When adding MT sliding to the previous mechanisms, we obtain the polarity profile *P*(*x*) from stochastic simulations for different sliding speeds *u*. All numerical values of model parameters are given in Table 2.

For the first and simplest scenario, we ignore MTs originating from the soma by setting 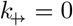, and let MTs whose plus ends reach the tip transition immediately to the shrinking state corresponding to the limit *k*_u_ → ∞. For this simple case, the polarity profile follows from Eq (72) as

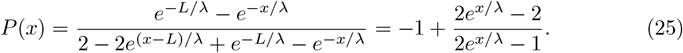

The polarity is negative along the entire neurite, see Fig 3A. The +OUT MTs are effectively restricted by the tip boundary, cannot grow further, and immediately transition to the shrinking state, while +IN MTs experience the open boundary at the base of the neurite; thereby, they have longer lifetimes and lengths compared to the +OUT MTs. At the neurite’s base, the polarity attains the value *P* (0) = − 1, as none of the +OUT MTs can cross the base because they nucleate within the neurite and extend towards the tip. Going from the base to the tip, the polarity becomes less negative with a characteristic length scale *λ*, reaching a maximum value at the tip *P*(*L*) = 1*/*(1 − 2*e*^*L/λ*^). For long neurites, *L* ≫ *λ*, the polarity near the tip of the neurite becomes zero, because of two effects compensating each other: +OUT MTs near the tip are depleted as they shrink upon contact with the tip, +IN MTs nucleate and point towards the soma such that their contribution to the local MT crossing numbers decays as we approach the neurite’s tip. It is important to note that even in this straightforward scenario in which MTs have identical dynamics regardless of their orientation, an asymmetric polarity emerges as a consequence of the spatial boundaries.

In scenario 2, we extend scenario 1 to include centrosomal MTs originating from the cell soma by using finite values for the rate 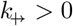 at which centrosomal MT cross the neurite’s base. From Eq (72), we obtain the polarity profile

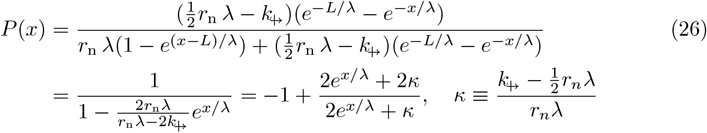

The above expression shows that when 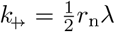 the polarity profile is zero along the whole neurite. In this case, MTs from the soma exactly compensate the absence of +OUT MTs near the base when the crossing rate is half of the nucleation rate within a length *λ*. Above this special value, 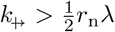 the polarity profile becomes entirely positive, while below it 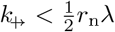 the polarity profile remains entirely negative, see Fig 3B.

For scenario 3, we additionally allow +OUT MTs to stay in the boundary state for a finite time by considering a finite unbinding rate *k*_u_. When the mean time in the boundary state is equal to the MT lifetime, 1*/k*_u_ = *τ*, we have an interesting case: a +OUT MT in the boundary state spends, on average, the same time it takes a growing +IN MT that exited the neurite at *x* = 0 to eventually return back by shrinking. In this scenario, MTs of the two orientations effectively experience the boundaries in the same way: for +OUT MTs, the tip of the neurite is effectively the same as the base of the neurite for +IN MTs. Therefore, the tip is enriched in +OUT MTs exactly in the same way as the base is enriched in +IN MTs. We recover this scenario from our analytical expression of the polarity profile from Eq (72) for 1*/k*_u_ = *τ*, and 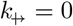, ignoring contributions from centrosomal MTs, as

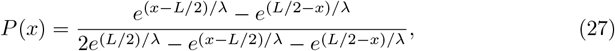

which is antisymmetric around the neurite’s center at *x* = *L/*2.

To further investigate scenario 3, we consider the numerical values given in Table 2 and determine the polarity profile from Eq (72) for different values of *k*_u_, see Fig 3C. As discussed before, when the residence time at the boundary state is equal to the lifetime of a free MT, 1*/k*_u_ ≈ *τ*, the base of the neurite displays a negative polarity that gradually increases to a positive polarity at the tip. For large residence times, 1*/k*_u_ ≫ *τ*, the +OUT MTs are highly stabilized at the tip and the overall polarity becomes positive. In the case of small residence times 1*/k*_u_ ≪ *τ*, the polarity profile is mostly negative with a positive region close to the tip. In any neurite with finite *k*_u_, the polarity is positive at the tip and exclusively +OUT oriented, since no +IN MTs can exist there as they could not have been nucleated from beyond the tip. It is important to note that the tip-stabilization mechanism favors +OUT MTs and can only lead to an axon-like MT organization in extreme tip stabilization *k*_u_ → 0, which would lead to an explosion in the number of +OUT MT, dominating +IN MTs.

Finally, for scenario 4, we include sliding at constant speed *u*. Recall that by construction in our model, when *u* is positive, the sliding mechanism carries +OUT MTs to the neurite’s tip and +IN MTs to the cell soma, and when *u* is negative, the other way around. To determine the polarity profile, we turn to stochastic simulations and take the ensemble average over 10000 simulations, as the equations are analytically intractable; see Event-driven simulations. For positive sliding speed *u*, sliding extends the positive polarity region near the tip because arriving +OUT MTs accumulate there, while newly nucleated +IN MTs are pushed away, see Fig 3D. Conversely, at the soma junction, it enhances the negative polarity by moving +OUT MTs soon after they appear away into the neurite. Sliding with negative speeds *u* reduces the number of +OUT MTs in the neurite, and it populates the tip with minus ends of +IN MTs, resulting in a negative polarity profile.

In summary, we explicitly show that the considered mechanisms can already bias the orientation of the MTs in opposite directions at the two ends of an otherwise featureless neurite. The orientational bias is the outcome of the competition of the mechanisms we introduced, with their magnitude determining the orientation bias favoring a +OUT/+IN bias as follows: more/less centrosomal MTs crossing into the neurite, slow/fast unbinding of boundary-stabilized +OUT MTs, and positive/negative sliding speed. The influence of these effects on polarity depends on the neurite length relative to the mean MT length *λ*: the shorter the neurite, the more prominent these mechanisms are, as they rely strongly on the effects induced by the boundaries.

So far, our model exploration shows that for moderate numerical values of parameters, we cannot obtain an organization with uniformly oriented MTs; an axonal organization with all +OUT MTs and a dendritic organization with all +IN MTs (for some species) is not possible since MTs of both orientations will always nucleate and be present to a degree. However, parallel nucleation, which we consider in the next subsection, allows us to get neurites with uniformly oriented MTs.

#### Parallel nucleation amplifies the polarity

In our framework, we propose that parallel amplification mechanisms enhance a polarity bias, either +IN or +OUT, contributing toward organizations of uniform orientation. Here, we study this effect through the parallel nucleation mechanism of our model. We consider two scenarios having the values of all parameters the same, differing only in the value of *k*_u_, such that when *K*_∥_ = 0, the first has a positive polarity across the neurite, which we identify as a +OUT bias, and the latter has a negative polarity across the neurite, which we identify as a +IN bias. We are interested in how the polarity profile behaves in the two scenarios as *K*_∥_ increases. The numerical values for the parameters of the two scenarios are presented in Table 3.

**Table 3.**
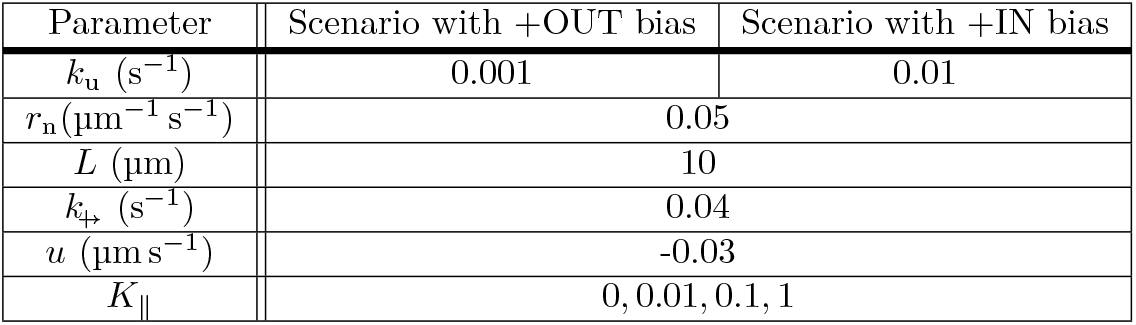
Numerical values of parameters for each of the +OUT and +IN bias scenarios considered in the study of the parallel nucleation mechanism and the results are obtained from stochastic simulations.

To study the effect of parallel nucleation on the two scenarios, we run simulations of the model as described in Event-driven simulations. In Fig 4 we present the polarity profiles for the two scenarios, where in panel A we have the scenario that is +OUT biased, and in panel B the scenario that is +IN biased, as their polarity profiles are entirely positive and entirely negative for the baseline case of *K*_∥_ = 0, respectively. In both scenarios, for increased *K*_∥_, the polarity profile is enhanced: the polarity of the +OUT scenario becomes more positive, and the polarity of the +IN scenario becomes more negative. At the highest value for *K*_∥_ = 1, the +OUT biased scenario attains an almost +OUT organization, i.e, *P*(*x*) = 1, while in the case of the +IN biased scenario the base remains mixed as the contribution of centrosomal +OUT MTs from the soma are unaffected by the nucleation mechanism.

**Fig 4.**
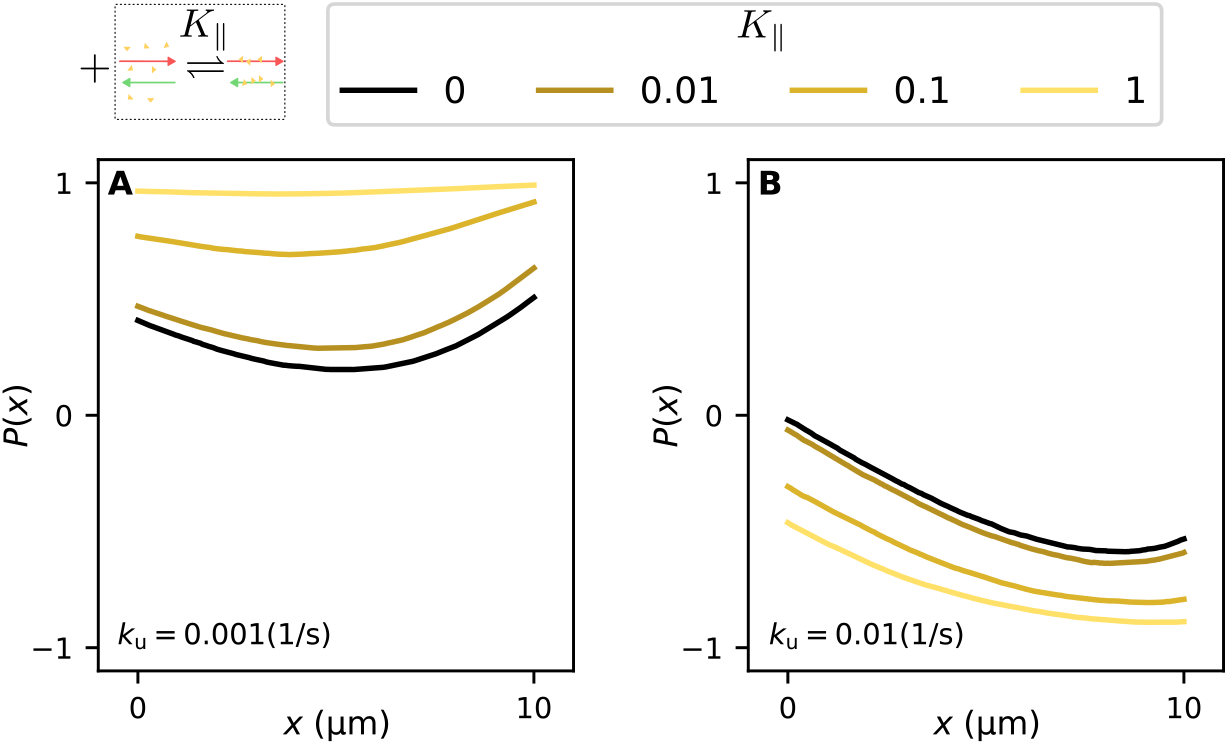
Parallel nucleation amplifies the polarity. We consider the effect of *K*_∥_ for two scenarios whose parameter sets differ only in *k*_u_ such that even for *K*_∥_ = 0 the polarity of the first is biased towards an axonal polarity and the polarity of the second is biased towards a dendritic one. (A) The polarity profile for *k*_u_ = 0.001 s^−1^ and in (B) the polarity profile for *k*_u_ = 0.01 s^−1^, for different values of *K*_∥_; the values for the remaining parameters are listed in Table 3

From this analysis, parallel nucleation appears consistent with the class of parallel amplification mechanisms we describe in our conceptual framework, see Conceptual framework: Three classes of mechanisms shaping the microtubule organization in developing neurons. We have shown that the unbinding rate *k*_u_ of MTs from the boundary state at the tip can serve as a control parameter for the bias, thereby governing axonal and dendritic organization. Integrating these results, in the next section, we introduce a description of a neuron as a system of neurites whose unbinding rates *k*_u_ are coupled through a shared polarity factor.

### Neuron model: Shared polarity and growth factors determine the microtubule organization in a developing neuron

We describe a developing neuron as a collection of *n* one-dimensional neurites connected to a reservoir representing the soma. From this reservoir, neurites exchange shared developmental factors that facilitate neurite growth and modulate MT dynamics, see Fig 5. In each neurite, MTs are subject to the same mechanisms discussed in the preceding sections; however, we now incorporate a dependence of the previously fixed parameters on the presence of factors in a neurite. We assume that motor proteins mediate the supply of these factors to the neurite; therefore, the number of polarity factors in a neurite is linked to its MT organization. This scenario falls within the class of polarization mechanisms in our conceptual framework, and it exhibits two major features: first, the MT organization and the number of factors in a neurite create a feedback loop; second, the MT organization and morphology are coupled across the neuron’s neurites.

**Fig 5.**
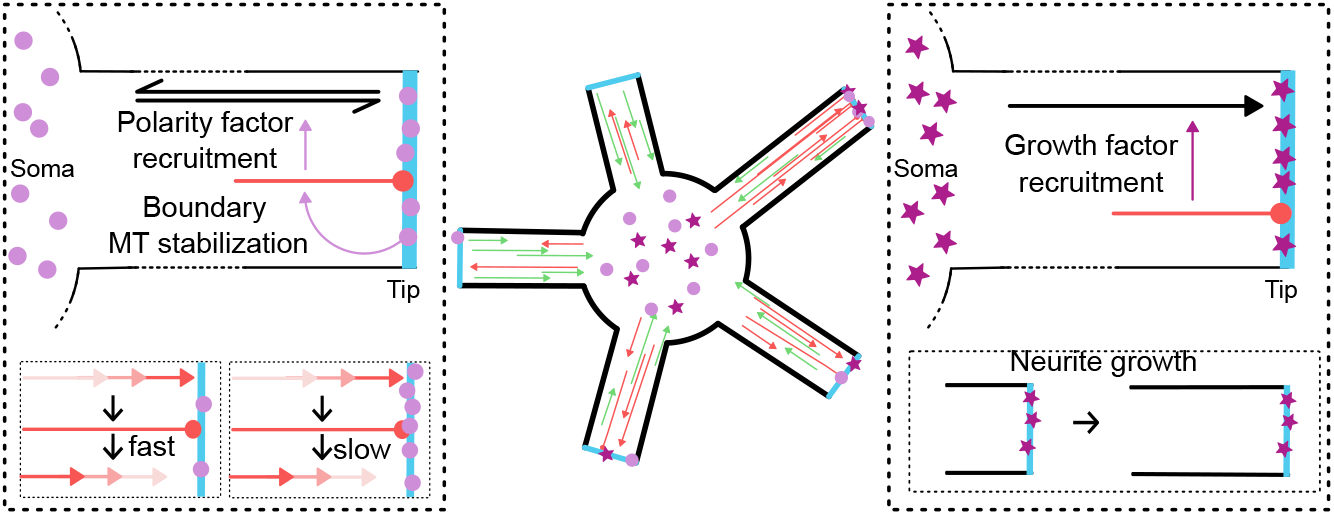
Shared polarity and growth factors interact with the MT dynamics in a neuron. (Center) We describe a neuron as a collection of neurites that are coupled via a shared pool of polarity and growth factors. (Left) polarity factors at the neurite’s tip are in a quasi-equilibrium with the somatic pool; the number of boundary MTs increases the recruitment of polarity factors, and the polarity factors increase the lifetime of MTs in the boundary state. (Right) The influx of growth factors at the neurite’s tip depends on the number of boundary MTs, and the elongation speed of the neurite is proportional to the number of growth factors.

During the early stages of neuronal development, the growth cone is a highly active center for the neurite’s fate and MT organization, motivating our following assumption: factor accumulation and influence on MT organization is predominantly on the growth cone of neurites, which in our model is the neurite tip. In particular, the transport of factors between neurites and soma is mass-action-like, with the limiting step in the influx being the number of +OUT bound MTs, which drives motor proteins to the growth cone. Moreover, for simplicity, we assume that the number of factors throughout the neuron is conserved, i.e., neither consumed nor produced. This last assumption keeps us from having additional parameters for the rates of factor production and consumption.

With the above assumptions, in quasi-steady state, we write the number *F*_*i*_ of a factor in neurite *i* as

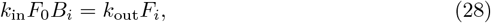

in which *k*_in_ and *k*_out_ are influx and outflux proportionality constants, *B*_*i*_ are the number of bound MTs in neurite *i*, and *F*_0_ is the number of factors in the soma.

Using the conservation of factors across the neuron, we write

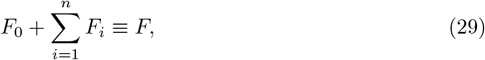

in which *F* is the total number of factors.

We then sum Eq (28) over all neurites and use Eq (29) to derive the number of factors left in the soma

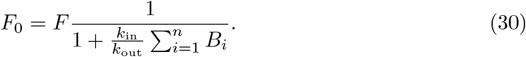

Combining Eq (30) and Eq (28), we obtain the number factors in each neurite

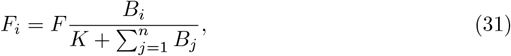

with the equilibrium constant 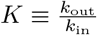.

Because the total number of a factor is conserved, the relevant quantity is the fraction of the factor in each neurite rather than its absolute number. Therefore, we introduce the normalized variables *f*_*i*_ ≡ *F*_*i*_*/F*.

In our model, we consider two types of shared developmental factors: (i) *Polarity factors*, which stabilize +OUT MTs bound at the tip, favoring a +OUT organization. (ii) *growth factors* whose presence in a neurite catalyzes the neurite’s elongation. The fraction of polarity factors *p*_*i*_ in neurite *i* follows from Eq (31) as

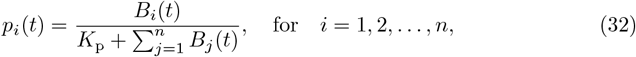

in which *K*_p_ is the polarity factor’s equilibrium constant. Similarly, the fraction of growth factors *g*_*i*_ for a neurite *i* is given by

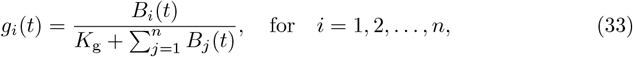

in which *K*_g_ is the growth factor’s equilibrium constant.

We consider that the polarity factors and the boundary MTs interact through a closed positive feedback loop by assuming that in a neurite, the transition rate *k*_u_ for the boundary MTs to the shrinking state depends on the number of polarity factors at the neurite’s tip, such that the more polarity factors there are, the longer the bound MTs stay bound. Mathematically, we introduce a dependence of *k*_u_ on the local *p* with a Hill function interpolating between two values, *k*_d_ and *k*_a_. Thus, we modify the rate *k*_u_ for neurite *i* to depend on the polarity factors as

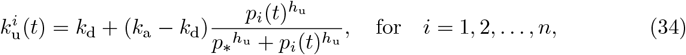

in which *h*_u_ is a Hill exponent and *p*_∗_ is the half-saturation fraction of polarity factors for which *k*_u_ is in the middle of the two values, (*k*_d_ + *k*_a_)*/*2. The above function assigns *k*_u_ a value between the baseline value *k*_d_ in the absence of the polarity factor and *k*_a_, corresponding to the case of *p*_*i*_ ≫ *p*_∗_. Consistent with the role of polarity factors in stabilizing boundary MTs, we consider *k*_a_ *< k*_d_.

The growth factor promotes the elongation of a neurite’s length *L*_*i*_ as

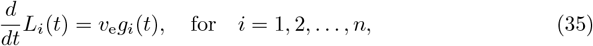

in which *v*_e_ is a proportionality speed, reflecting the maximum elongation speed a neurite can achieve when it takes over all growth factors.

These equations complete our description of a neuron as a coupled system of several neurites that exchange polarity and growth factors. In the next section, we will investigate the consequences of these mechanisms on the morphology of a neuron.

#### The multi-neurite model recapitulates physiological neuronal development

To study the development of the MT organization in neurons, we simulate our neuron model as described in the previous section. We consider neurons with 5 neurites, initially without MTs and a length of *L* = 2 µm. We run simulations as specified in Event-driven simulations, using parameters listed in Table 4 to determine the length and mean polarity of each neurite.

**Table 4.** Numerical values for parameters used for the simulations of the neuron model.

| Parameter | Values |
| --- | --- |
| $p_*$ | 0.2, 0.4, 0.6 |
| $k_a$ ($\text{s}^{-1}$) | 0.001 |
| $k_d$ ($\text{s}^{-1}$) | 0.01 |
| $h_u$ | 5 |
| $K_p$ | 10 |
| $K_g$ | 0.1 |
| $K_{\parallel}$ | 1 |
| $r_n$ ($\mu\text{m}^{-1} \text{s}^{-1}$) | 0.05 |
| $L_{t=0}$ ($\mu\text{m}$) | 2 |
| $k_{\leftrightarrow}$ ($\text{s}^{-1}$) | 0.04 |
| $u$ ($\mu\text{m} \text{s}^{-1}$) | -0.03 |

We are particularly interested in how the polarization mechanism determines the mean polarity and length of a neuron’s neurites. We study the model for three values of the half-saturation parameter *p*_∗_ : 0.2, 0.4, and 0.6. For each scenario, we simulate the first 24 hours of development for 100 neurons and record the mean polarity and length for each neurite. Because the mean polarity fluctuates significantly due to small-number effects in the MT populations, we define the representative end-of-simulation polarity as the time average of samples taken every minute during the final hour. Fig 6A - C shows the joint distribution of lengths and mean polarities at the end of the simulation. Across these scenarios, we observe that the distribution of mean polarities is unimodal for *p*_∗_ = 0.6 with a peak at negative values, and bimodal for *p*_∗_ = 0.2 and *p*_∗_ = 0.4 with one peak at negative values and a second peak around 1. Based on this bimodality, we set a threshold at ⟨*P*⟩ = 0.75 to classify neurites with ⟨*P*⟩ ≥ 0.75 into axons, and neurites with ⟨*P*⟩ *<* 0.75 into dendrites.

**Fig 6.**
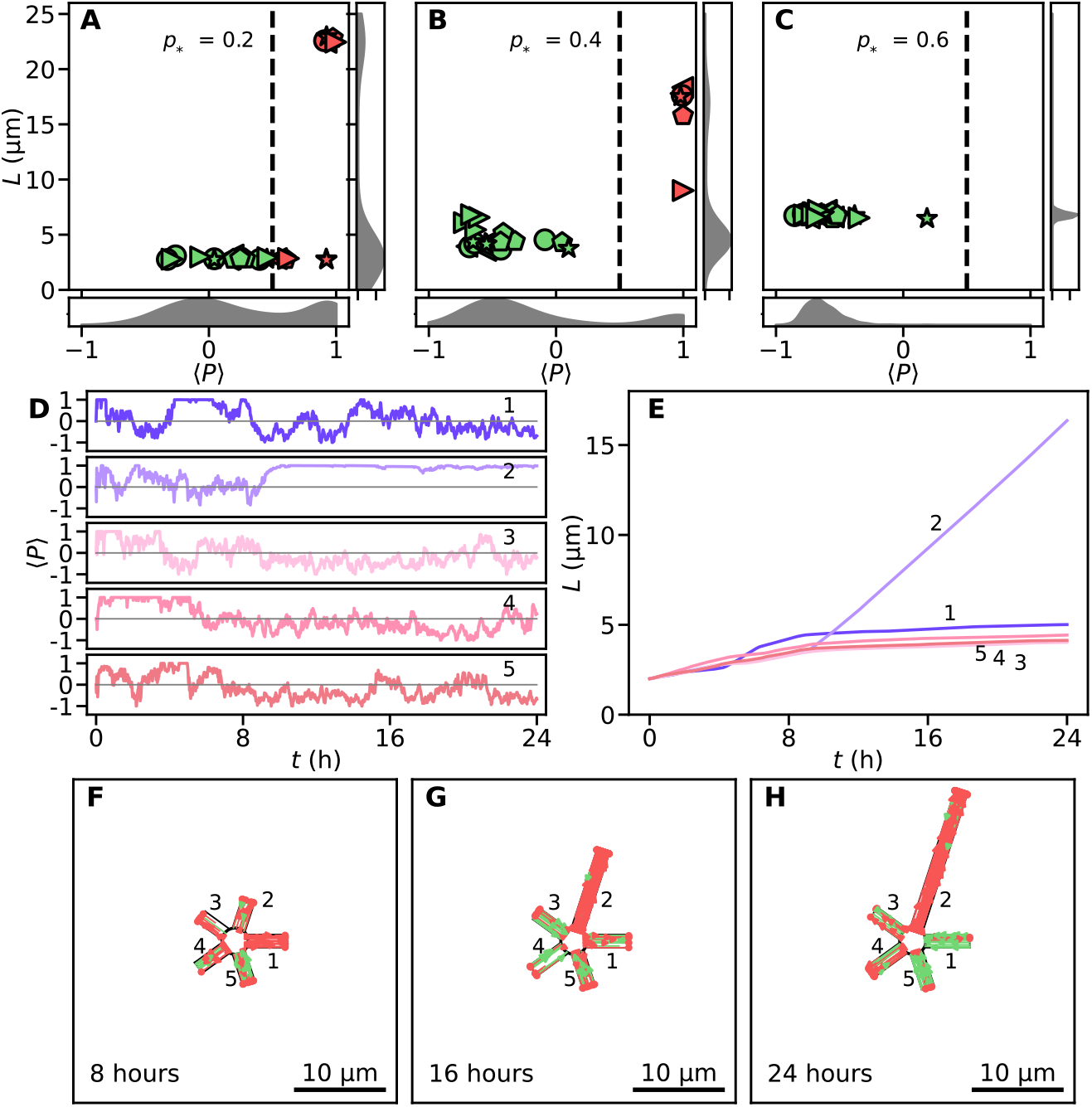
Neuronal polarization induced by the competition of shared polarity and growth factors. We simulate neurons with numerical values for the parameters listed in Table 4. For each condition, we report results from 100 independent runs. (A-C): Estimates of densities (gray) of the mean polarity and neurites’ length for different values of the half-saturation parameter *p*_∗_ (A): 0.2, (B): 0.4, and (C): 0.6. This parameter determines how the lifetime of MTs in the boundary state depends on the number of polarity factors. We highlight the neurites of five selected neurons with individual scatter markers to show their mean polarities and lengths. Each marker represents a neurite with axon-like (green) or dendritic-like (red) identities. These identities are based on a threshold of ⟨*P*⟩ = 0.75 (dashed vertical line), chosen to separate the peaks of the mean polarity density. Neurons with mostly single axons appear for a moderate value of *p*_∗_ = 0.4 in (B). (D) The evolution of the mean polarities in time of 5 neurites (enumerated) for a realization for *p*_∗_ = 0.4, and (E) the time series of their length. While initially neurite 1 gains a high polarity between 5 and 7 hours, neurite 2 becomes the axon at around 8 hours accompanied by faster and longer elongation, see (D) and (E). (F-H) Illustrates the MT organization in the simulated developing neuron corresponding to panels D and E, from left to right, snapshots at 8, 16, and 24 hours. Red indicates +OUT MTs and green +IN MTs.

For *p*_∗_ = 0.2, many neurons develop more than one axon; in particular, of the 100 neurons, 2, 62, 31, and 5 developed 0, 1, 2, 3, and 4 axons, respectively. Neurites classified as axons have, on average, a length of around 22.5 µm, while dendritic neurites have, on average, a length of 2.8 µm. We observe a positive correlation between mean polarity and neurite length, as expected: higher polarity corresponds to a larger population of boundary MTs, which in turn redirect more growth factor and yield faster neurite elongation. Essentially, many of the neurons in this scenario develop multiple axons because *p*_∗_ is low enough that multiple neurites could reach the axonal level without significant competition.

For *p*_∗_ = 0.4, we find that almost all neurons develop a single axon; in particular, of the 100 neurons, 6, 92, and 2 developed 0, 1, and 2 axons, respectively. Most neurites with an axonal fate have a length of around 16 µm, while dendritic neurites have a length of 5 µm. The length distributions of both dendrites and axons are slightly broader than in the other scenarios because the neuron occasionally switches the designated neurite with the axonal fate, something we discuss further later. This scenario is consistent with the physiological development of neurons, i.e., starting with multiple equivalent neurites, one emerges as the axon with a uniform +OUT polarity, while the others have mixed or negative polarity, and the axon is significantly longer than the other neurites. Moreover, under these conditions, we capture the transient exchange of axonal fate as suggested in experiments [16, 58–60].

For *p*_∗_ = 0.6, we find that none of the neurons develop any axon, and neurites have an identical length of around 6 µm. In this case, the average number of bound MT that the axonal and dendritic fates can support is not enough to consistently recruit enough polarity factors above *p*_∗_= 0.6. Occasionally, by number fluctuations, enough bound MT are present for some neurites to briefly transition from the dendritic fate to the axonal fate. However, this state is not sustained for long, as small number fluctuations let neurites that reach the axonal level drop back to the dendritic level. In this case, all neurites get to be in the axonal state for a short while, and therefore they grow more than the corresponding dendrites from scenarios with *p*_∗_ = 0.2 and *p*_∗_ = 0.4, in which one or more neurites take the polarity factors limiting them from the other neurites.

For a representative neuron with *p*_∗_ = 0.4, which is consistent with the physiological development, we determine the time series of the mean polarities and lengths of its neurites over the first 24 hours of development, see Fig 6D and E. In the first few hours, we observe multiple neurites with a positive mean polarity, some reaching a polarity of 1, see Fig 6D. This early axon-like organization for multiple neurites has also been observed experimentally [61, 62]. At this stage of development, all neurites that start from a length of 2 µm grow to around 5 µm, which is about half the mean MT length *λ*. The centrosomal +OUT MTs from the soma dominate the MT population, and their population is further reinforced by parallel nucleation. In other words, at these neurite lengths, there is a net +OUT bias which is enhanced by parallel nucleation so that multiple neurites adopt an axonal fate. At this stage, multiple neurites can become axon-like, even without the effect of the polarity factor. At around 5 hours, as neurites have elongated sufficiently, the non-centrosomal MTs nucleated in the neurite outnumber the centrosomal MTs and a single neurite, neurite 1, adopts a temporary axonal fate, see Fig 6D. About three hours later, neurite 1 changes to a dendrite fate, and neurite 2 takes its place as the axon, in which it stays until the end of the simulation. This switch occurs because of small-number fluctuations of MTs. Neurite 2 remains an axon for the rest of the simulation, while the other neurites stay in the dendritic fate. Neurite 2 also outgrows the other neurites in length as it accumulates the growth factors, see Fig 6E. In the simulation of this neuron, the MT organization in the neurites transiently changes until one neurite significantly outgrows and adopts an axon-like MT organization with mostly +OUT orientation, see Fig 6F-H.

These findings are consistent with the local mechanisms that we identified and discussed in the model of a single neurite. Therefore, we further substantiate our findings with analytical considerations for a simplified description. Although neurites continuously elongate and the whole system does not settle into a steady state, the elongation timescale on the order of µm h^−1^ is much slower than MT turnover on µm s^−1^. It is therefore natural to view the organization as a quasi-steady state with slowly changing neurite lengths. In this regime, and for an abrupt change of *k*_u_ in response to the polarity factor *p*, the transition rate *k*_u_ effectively takes one of two levels depending on whether the local polarity factor surpasses the threshold or not. These two levels correspond to two distinct steady populations of boundary MTs and, therefore, to two distinct polarity levels, which we consider in this limit as the axon level and the dendrite level. For a bifurcation analysis on the number of neurites in each fate/level, see Bifurcation analysis of the neuron model.

## Discussion

We summarize the intuition derived from our model as an interplay of three key mechanisms; (i) The asymmetric geometry of neurites, as elongated compartments with a closed tip and an open connection to the soma, biases how mechanisms act on +OUT and +IN MTs, with the characteristic length scale of these boundary effects set by the mean length of MTs. (ii) Bias mechanisms alone cannot generate the orientation of an axon with complete +OUT nor, in the case of *Drosophila*, of dendrites with complete +IN [5, 6]. Therefore, they must be complemented by amplification mechanisms such as biased nucleation [19, 20, 55], bundling [11, 23], and severing [63] to establish the proper microtubule organization in neurons. (iii) Considering the polarization process, we show that under sufficient conditions, proper neuronal development can emerge from resource competition. Our biophysical model reveals a novel explanation for the oscillating identity of the axon-like neurite observed in developing neurons, both in experiments [16, 58–60] and in our model: fluctuations in the number of MTs within a neurite can change which neurite is selected as the axon, as the polarity mechanism switches between them. This is the simplest explanation proposed so far and requires no additional biochemical or biophysical machinery, although some underlying feedback is presumably still at play.

Although our models do not explicitly include the large set of molecular components proposed in the literature, these components are, in a sense, already present in a coarse-grained form–effectively averaged into the model and its parameters. Our proposed framework can nonetheless be used to extrapolate how a given component or mechanism would affect the overall organization. Thus, a natural next step is to incorporate specific components into the concrete mechanisms that constitute them to investigate their biophysical detail, while keeping in mind that, as our model shows, such mechanisms can interact in ways the coarse-grained framework cannot capture.

Several components are good candidates for such detailed refinements. The growth cone, which we represent simply as the boundary at the neurite tip, in reality hosts multiple molecular players that could carry out the stabilization mechanism, including the actin network and its associated proteins [64, 65]. The polarization mechanism, treated here agnostically, likewise needs to be resolved into an interplay of motors, actin, and other factors that together create a winner-takes-all mechanism [38, 59]. Finally and importantly, our model treats all MTs as a single subset, with their dynamics effectively averaged; a more complete description must distinguish the different MT subsets within a neurite and how they inter-convert, since many mechanisms act selectively on specific subsets [12, 66].

It is worth noting that in our description, the influence of the geometric bias on polarity diminishes as neurites elongate, see, for example, the large-*L* results in Fig 3. While this vanishing bias could be interpreted as a weakness of the model, it points to an important feature of long-term neuronal polarization: the MT organization must have memory. Such memory could be provided by mechanisms that preserve MT orientation over long time scales in very long neurites, and it connects to the well-known argument that neuronal MTs, just like in other cells, comprise distinct subsets [67]. One classification separates dynamic from stable MTs [66]: dynamic MTs undergo rapid dynamic instability of growth and shrinkage, whereas stable MTs are long-lived, maintained by MT-associated proteins, post-translational modifications, capping, and bundling. We therefore hypothesize that during early development, the rapidly changing organization is primarily driven by dynamic MTs, which gradually stabilize as development proceeds. In this way, a neurite can retain its distinct polarity type even after substantial growth that would otherwise diminish the original biasing mechanisms.

Within the broader context of neuronal development, we addressed questions specific to the organization of the MT cytoskeleton; however, our findings remain confined to this subsystem. Future work must establish explicit connections to the other cytoskeletal components and the broader mechanisms underlying neuronal development, the establishment of axonal and dendritic identity, and, ultimately, the formation of functional neural networks.

## Materials and Methods

### Parallel nucleation kinetics

We consider MT nucleators that can be either free or bound to MTs. Free nucleators nucleate MTs with equally likely orientations, whereas nucleators bound to a MT nucleate new filaments with the same orientation as the one they are bound to. For simplicity, we assume that bound and free nucleators nucleate MTs spontaneously with the same rate *k*_n_. We introduce the orientation-resolved nucleation rate densities, 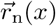 and 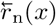, as

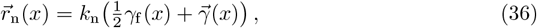

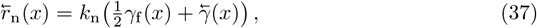

in which *k*_n_ is the nucleation rate, *γ*_f_(*x*) is the linear density of free nucleators, and 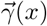 and 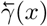 the densities of nucleators bound to +OUT MTs and +IN MTs, respectively.

We assume that the total number of nucleators in a neurite of length *L* is constant and equal to 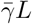, where 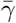 is a proportionality constant with units of linear density.

Moreover, nucleators diffuse rapidly and bind to or unbind from MTs on a timescale much faster than the MT turnover. Consequently, the free nucleator density is uniform, *γ*_f_(*x*) = *γ*_f_, and the bound nucleators follow the local equilibrium relations:

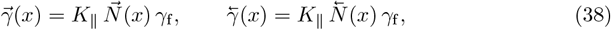

in which *K*_∥_ is the ratio of the binding and unbinding rates, and 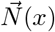 and 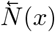 are the crossing numbers of MTs with +OUT and +IN orientations, respectively.

Conservation of nucleators requires that the total number of free and bound nucleators equals 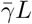, therefore

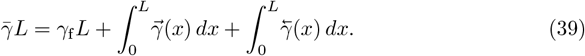

By substituting the equilibrium relations Eq (38) and carrying out the integrals, we obtain

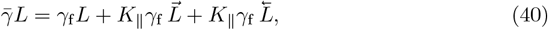

with 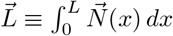 and 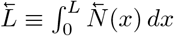 the total lengths of MTs corresponding to the two orientations. Solving Eq (40) with respect to *γ*_f_ we find

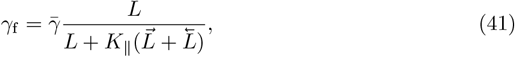

which we substitute in Eq (38) to get

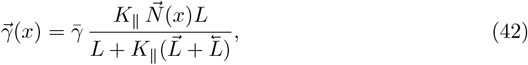

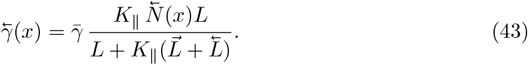

Using the expressions for 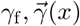 and 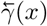 in Eq (36)–Eq (37) we obtain the explicit solutions of the nucleation rate densities,

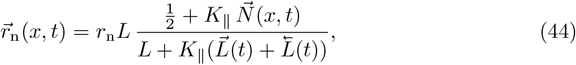

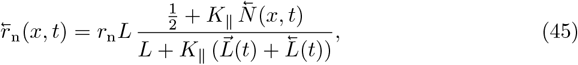

in which 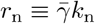 is the nucleation rate per unit length.

Finally, by integrating 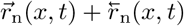 along the neurite we find

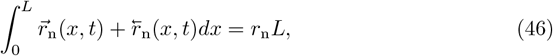

confirming that *r*_n_ indeed represents the total nucleation rate density.

### Derivation of analytical solutions

For a simplified scenario, we derive analytical expressions for the MT densities, local MT crossing numbers, and the polarity profile of the neurite model presented in A biophysical model of microtubule organization in a single neurite at steady state. We consider a neurite of length *L*, and ignore MT sliding by setting *u* = 0. Moreover, we assume that all nucleators are free, i.e., *K*_∥_ = 0, so that a nucleation event nucleates MTs equally likely in both orientations, 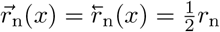.

Starting with the neurite nucleated MTs, under our simplifying assumptions the dynamical equations Eq (7)-Eq (10) at steady-state read

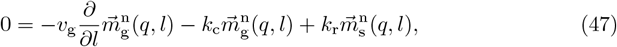

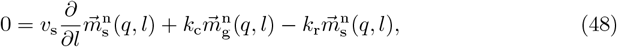

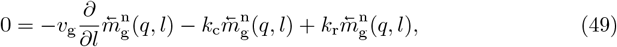

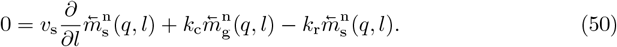

The boundary dynamics in Eq (11) at steady state and the corresponding flux balance in Eq (12) simplify to

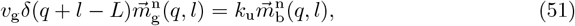

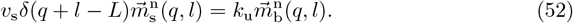

The nucleation flux balance from Eq (19) for *K*_∥_ = 0 reads

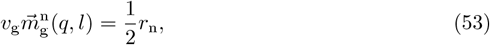

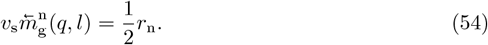

The pairs of differential equations Eq (47)-Eq (48) and Eq (49)-Eq (50) correspond to the classic Dogterom–Leibler steady-state relations for dynamic instability [57]. In our case, however, the minus ends of neurite-nucleated MTs are uniformly distributed along the neurite domain [0, *L*]. The solutions for the growing and shrinking subpopulations read

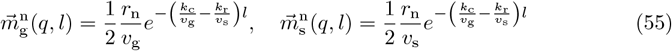

for MTs in the +OUT orientation, which are identical to the corresponding densities for MTs in the +IN orientation

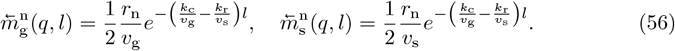

Notice that the tip boundary does not affect the growing and shrinking subpopulations of +OUT MTs, since the flux-balance condition 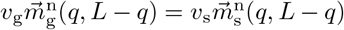, implied by Eq (51), is already satisfied by 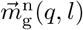 and 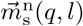. We obtain the density of MTs in the boundary state by substituting 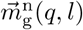 in Eq (51) as

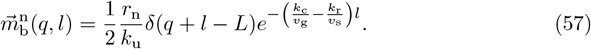

Adding up the densities over all plus-end states, we determine the densities of the non-centrosomal microtubules of the two orientations,

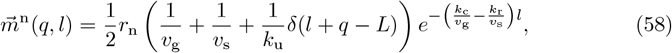

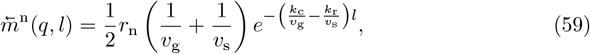

The differential equations Eq (14) and Eq (15) for the growing and shrinking centrosomal MTs that originate from the soma read

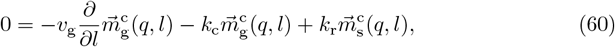

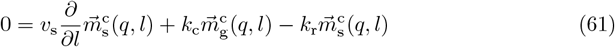

with the nucleation condition,

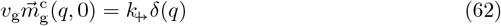

and boundary conditions

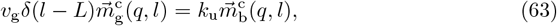

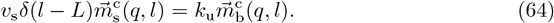

We solve the above equations and sum over all plus-end states to find that

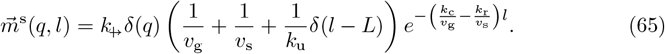

In the original Dogterom–Leibler description [57], the effect of the dynamic instability parameters on the steady-state MT-length distribution can be reduced to two effective parameters: the mean MT length

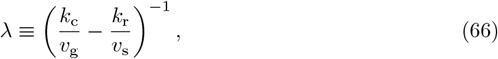

and the MT lifetime

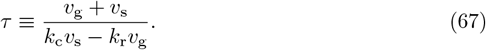

Therefore, we rewrite our MT densities using *λ* and *τ*, which simplifies the effects of dynamic instability to

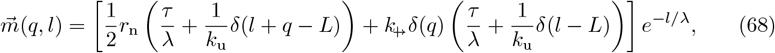

and

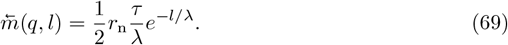

Next, we derive the local MT crossing numbers at position *x* by integrating 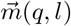 and 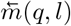 as in Eq (5) and Eq (6) respectively. The local MT crossing numbers for the two orientations read

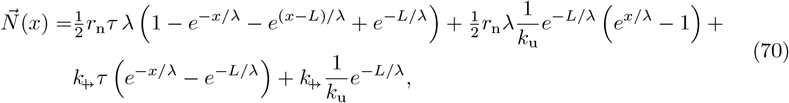

and

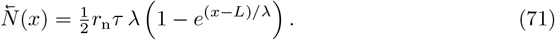

Using 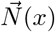 and 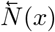, we find an analytical expression for the polarity profile defined in Eq (1) as

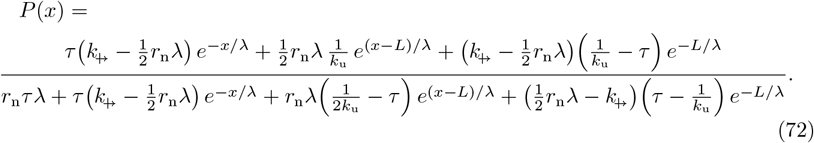

The mean polarity ⟨*P*⟩ from Eq (2) does not have a simple closed form. However, it is straightforward to evaluate it by numerical integration of *P*(*x*).

### Derivation of semi-analytical solutions for non-zero *K*_∥_

We generalize the analysis from Derivation of analytical solutions for the case where *K*_∥_ ≠ 0. In this more general case, the length dynamics are unchanged, so the *l*-dependence of the *m*-densities remains the same, while the minus ends are distributed according to the nucleation conditions Eq (18). Therefore, we can reuse the expressions for the MT densities from the last analysis, but keep the general nucleation condition Eq (18) instead of 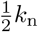,

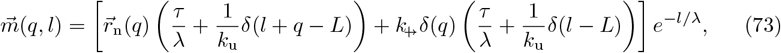

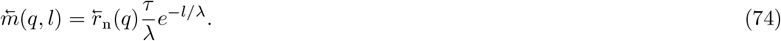

Since 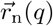 and 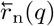 depend on 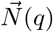 and 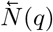, see Eq (18), we seek analytical self-consistent solutions to Eq (73) and Eq (74). Integrating both sides of Eq (73) and Eq (74) as in Eq (5) and Eq (6) we get

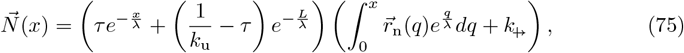

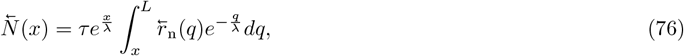

which upon substitution of the nucleation conditions Eq (18) and rewriting using the dimensionless parameters introduced in Derivation of analytical solutions, lead to the following integral equations

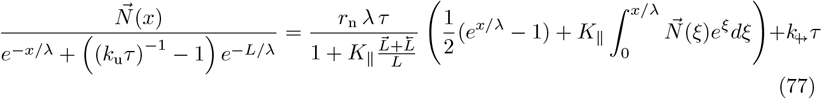

and

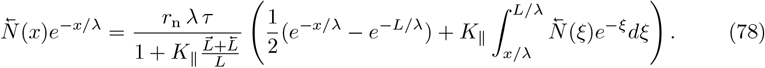

By taking a derivative with respect to *x* on both sides of the above integral equations, we convert them to first-order differential equations, which can then be solved using integrating factors as

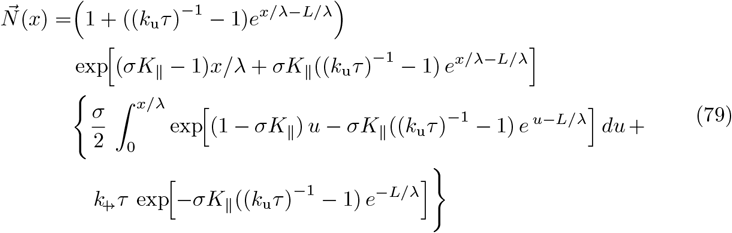

and

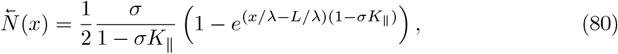

in which 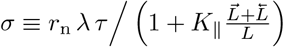.

From the expressions Eq (79) and Eq (80) we see that 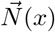 and 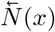 parametrically depend on 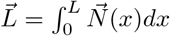 and 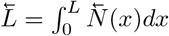 through *σ*. For any given parameter set, we determine *σ* using the bisection method: in each iteration, we numerically integrate the right-hand sides of Eq (79) and Eq (80) to obtain 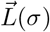 and 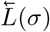 until the self-consistency condition is satisfied to sufficient tolerance.

This is as far as we can proceed analytically with the neurite model; we therefore cross-validate these semi-analytical results against simulations in Event-driven simulations.

### Event-driven simulations

We construct an exact, event-driven algorithm for the stochastic processes that describe the dynamics of neurites and their MT populations, corresponding to our model in A biophysical model of microtubule organization in a single neurite and Neuron model: Shared polarity and growth factors determine the microtubule organization in a developing neuron. In the following, we present the algorithm for a neuron with several neurites; the case of a single neurite with fixed length is a special case of this general setting. The implementation, together with documentation, can be found on the following remote repository https://codeberg.org/kyriacos/microtubule-organization-neurons.

Each iteration of the simulation consists of the following steps: (i) sample the waiting time Δ*t* to the next event; (ii) update the neurite lengths and the lengths and minus-end positions of all MTs up to time *t* + Δ*t*; (iii) execute the event; and (iv) repeat until the desired simulation time is reached.

In our model, we distinguish stochastic and deterministic events. The stochastic events are catastrophe, rescue, nucleation, entry of MTs from the soma, and exit from the boundary state. The deterministic events are: (i) when a shrinking MT reaches zero length and disappears, and (ii) when a growing MT reaches the neurite tip and transitions into the boundary state.

Events of catastrophe, rescue, soma entry, and leaving the boundary state have constant propensities at each time step, so we sample the earliest of these events using the standard Gillespie procedure [68]. We refer to these as *simple* stochastic events. The waiting time to the next such event is therefore

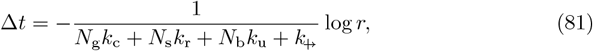

in which *N*_g_, *N*_s_, and *N*_b_ are the total numbers of growing, shrinking, and boundary MTs, respectively, and *r* is a random number uniformly sampled from [0, 1]. The type of event is then drawn proportionally to its propensity, e.g. *N*_g_*k*_c_ for a catastrophe event. For a sampled catastrophe, rescue, or unbinding event, a MT is sampled equally likely from the corresponding subpopulation, as they contribute equally to the propensity, while for a sampled crossing event, a new MT is considered.

To sample nucleation events in a neurite, we need to consider that the nucleation rate densities Eq (18) are spatially and temporally dependent propensities. For notational clarity, we consider a single neurite and drop the index *i*. The functions 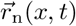 and 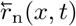 depend on the MT configuration through the local MT crossing numbers 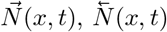 and the total MT lengths 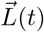 and 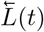 hence they are spatially heterogeneous and time dependent.

A convenient procedure is to first sample the waiting time Δ*t* to the next nucleation event from the *total* nucleation rate *r*_n_*L*(*t*), and then, conditional on that time, sample the nucleation position from the total spatial nucleation density at the propagated time, 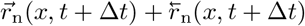, and finally sample the nucleation orientation.

At any time *t* the total nucleation rate is the integral of the orientation-resolved nucleation densities, which evaluates to *r*_n_*L*(*t*), see Eq (46). Within a timestep, the number of boundary MTs remains constant, and thus the neurite grows at constant speed according to Eq (35)

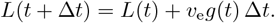

Therefore, the instantaneous total nucleation propensity is

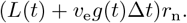

The survival probability for having no nucleation during a time interval Δ*t* is

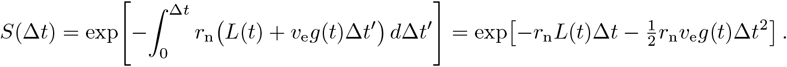

Using inverse-transform sampling, the waiting time for nucleation is

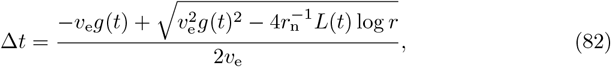

where *r* is a uniformly distributed random number in [0, 1].

After sampling Δ*t*, we determine the position of the candidate nucleation event. The total spatial nucleation density is proportional to

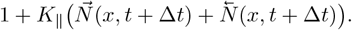

We can draw positions by rejection sampling: a candidate location is drawn uniformly from [0, *L*(*t* + Δ*t*)] and accepted with probability

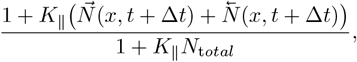

where *N*_t*otal*_ is the total number of MTs in the neurite and therefore an upper bound for the numerator.

Finally, given the accepted position *x*^*′*^, the orientation for the nucleating MTs is chosen with probabilities

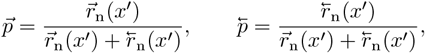

for the +OUT and +IN orientations, respectively.

We now consider the time intervals associated with deterministic events. A shrinking MT of length *l*(*t*) disappears when its length reaches zero, which occurs after

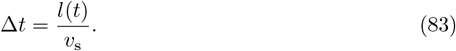

Similarly, a growing MT reaches the neurite tip when *q* + *l* = *L*(*t*). Since the tip moves with speed *v*_e_*g*(*t*), the relative speed of the plus end with respect to the tip is *v*_g_ + *u* − *v*_e_*g*(*t*), giving the waiting time

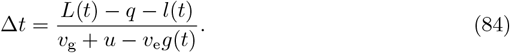

For each event type, stochastic or deterministic, we compute a candidate waiting time Δ*t*. The next event is determined by the smallest of these times. Before executing that event, we propagate the system deterministically to time *t* + Δ*t*. The MT lengths evolve as

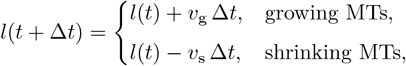

and for all neurite-nucleated MTs that are not in the boundary state, we update the minus-end positions according to sliding:

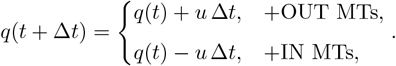

Finally, we execute the chosen event: nucleation introduces a new MT of zero length; a catastrophe, rescue, or boundary-exit changes the dynamical state of the selected MT; and when a MT reaches zero length, it is removed from the system. In the whole-neuron simulations, the numbers of growth factor and polarity factor in each neurite are updated according to the equations in Neuron model: Shared polarity and growth factors determine the microtubule organization in a developing neuron, given that after the execution of the event, the number of bound MTs could have changed. The unbinding rate *k*_u_ is then updated based on the new fraction of polarity factors, and the neurite length evolves as

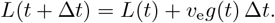

To validate the correspondence between the density-based description of the main text and the event-driven implementation, we compare the simulations to the semi-analytical solution in Derivation of semi-analytical solutions for non-zero *K*_∥_; see Fig 7. The agreement between the semi-analytical curves and the ensemble averages of the stochastic simulations is excellent.

**Fig 7.**
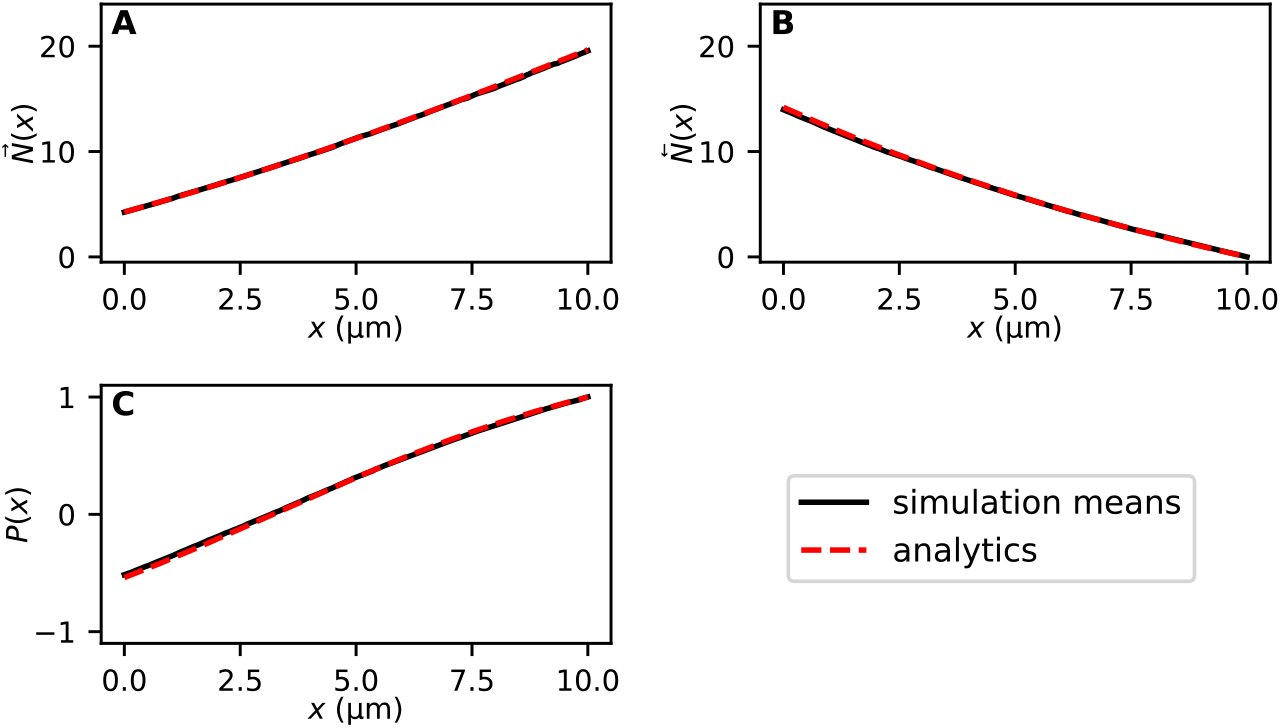
Validation of the stochastic simulator with semi-analytical solutions. (A and B) The local MT crossing numbers for +OUT and +IN MTs, (C) the corresponding polarity profile. The black, solid lines are simulation results, and the red, dashed lines are the solutions from Derivation of semi-analytical solutions for non-zero *K*_∥_. We use the following numerical values for the parameters: *L* = 10 µm, *r*_n_ = 0.05 µm^−1^ s^−1^, *k*_c_ = 0.12 s^−1^, *k*_r_ = 0.29 s^−1^, *v*_g_ = 0.10 µm s^−1^, *v*_s_ = 0.27 µm s^−1^, *k*_u_ = 0.01 s^−1^, 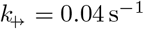, *K*_∥_ = 0.1. We simulated the time evolution of 1000 independent systems for 1 *×* 10^5^ s from which we sampled the MT configuration at the end of the simulation and determined the ensemble averages.

### Numerical values for model parameters

Our model contains parameters describing MT dynamic instability, nucleation, sliding, transition into and out of the boundary state, and neurite elongation. Whenever possible, we selected values based on experimental measurements in developing neurons; otherwise, we chose values guided by analytical calculations to ensure that the resulting MT numbers and length scales fall within biologically realistic ranges. Table 5 summarizes a range of reasonable values for our model parameters.

**Table 5.**
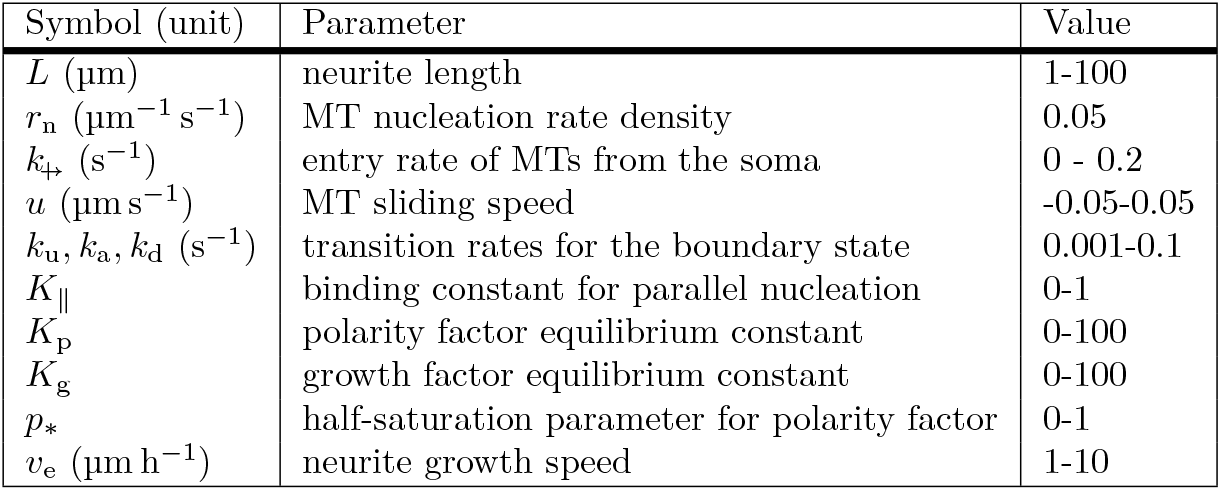
Model parameters and reasonable ranges of numerical values. These values are based on analysis assuming the dynamic instability parameters shown in Table 1

| Symbol (unit) | Parameter | Value |
| --- | --- | --- |
| $L$ ($\mu\text{m}$) | neurite length | 1-100 |
| $r_n$ ($\mu\text{m}^{-1} \text{s}^{-1}$) | MT nucleation rate density | 0.05 |
| $k_{\text{in}}$ ($\text{s}^{-1}$) | entry rate of MTs from the soma | 0 - 0.2 |
| $u$ ($\mu\text{m} \text{s}^{-1}$) | MT sliding speed | -0.05-0.05 |
| $k_u, k_a, k_d$ ($\text{s}^{-1}$) | transition rates for the boundary state | 0.001-0.1 |
| $K_{\parallel}$ | binding constant for parallel nucleation | 0-1 |
| $K_p$ | polarity factor equilibrium constant | 0-100 |
| $K_g$ | growth factor equilibrium constant | 0-100 |
| $p_*$ | half-saturation parameter for polarity factor | 0-1 |
| $v_e$ ($\mu\text{m} \text{h}^{-1}$) | neurite growth speed | 1-10 |

#### Dynamic instability

We use catastrophe and rescue rates *k*_c_ = 0.12 s^−1^ and *k*_r_ = 0.29 s^−1^, and growth and shrinkage speeds *v*_g_ = 0.1 µm s^−1^ and *v*_s_ = 0.27 µm s^−1^, measured in developing *Caenorhabditis elegans* DA9 motor neurons [69], which is one of the few datasets reporting all four parameters in a developing neuron. These values yield a mean MT length, calculated using Eq (23), of *λ* = 7.9 µm, consistent with measurements in young neurites [69–71], and a mean MT lifetime, calculated from Eq (24), of *τ* = 109 s.

#### Nucleation rate density

To set the nucleation rate density *r*_n_, we use a simple estimate based on the expected number of MTs crossing a typical neurite cross-section, which is within 10-100 MTs [72]. In the absence of spatial boundaries and sliding, MTs nucleate uniformly along the neurite with rate density *r*_n_ and subsequently undergo dynamic instability. The nucleation of MTs and their catastrophe back to zero length defines a standard birth–death process: minus ends appear at rate *r*_n_ per unit length and survive for an average time equal to the mean MT lifetime *τ*. Thus, the resulting linear density of minus ends is *r*_n_ *τ*. To estimate the mean number of MTs crossing any position *x* along the neurite, note that a MT of mean length *λ* covers, on average, an interval of size *λ*, so the mean number of MTs at a cross-section is *r*_n_ *τ λ*. For *τ* = 109 s and *λ* = 7.9 µm, derived from the set of dynamic instability parameters, choosing *r*_n_ = 0.05 µm^−1^ s^−1^ yields 43 MTs which is consistent with literature [72, 73].

#### Centrosomal MTs originating in the soma

In our description, centrosomal MTs enter the neurite at a rate 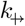. As for non-centrosomal MTs in the neurite, a centrosomal MT from the cell soma persists on average for a time *τ*, so their mean number in a neurite is 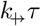.

At the base of the neurite, we require that non-centrosomal MTs from the neurite do not dominate the local MT population. Because non-centrosomal MTs from the neurite comprise on average 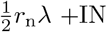 +IN filaments crossing any given position, we impose 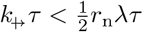, which gives the upper bound 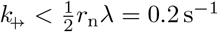. In practice, we use values far below this bound; throughout the simulations 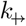 is chosen such that centrosomal MTs from the cell soma contribute roughly 10% of the total MT numbers at the base.

#### Motor-driven sliding speed

Live-cell imaging of MTs in neurons suggests numerical values for the speed of MT sliding that spans 3 orders of magnitude, ranging from 0.001 µm s^−1^ [16, 61] to 1 µm s^−1^ [74]. We consider the geometric mean of the above values which is equal to 0.05 µm s^−1^ as a representative upper bound on the sliding speed.

#### Transition rate from the boundary state

We consider that the mean time at the boundary state, which is the inverse of the unbinding rate, should be within an order of magnitude above and below the MT lifetime, therefore, 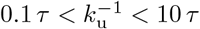, which corresponds to a range for *k*_u_ from 0.1 s^−1^ to 0.001 s^−1^.

#### Parallel-nucleation

The binding constant *K*_∥_ modulates the ratio of bound to free nucleating factors. This fraction can be obtained by adding and integrating equations Eq (38) to

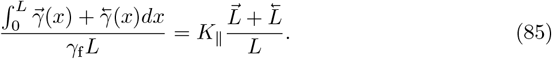

The total accumulated length of all MTs in a neurite 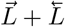 is typically on the order of tens of mean MT lengths in a young neurite of length *L* ~ *λ*. For example, for a neurite with length *L* = *λ* containing around 50 MTs and with *K*_∥_ = 1, 98% of nucleators are bound. Therefore, values of *K*_∥_ *>* 1 do not make much difference, so we consider [0, 1] as a reasonable range for *K*_∥_.

#### Extension speed of the neurite

Axons in young neurons have been reported to grow tens of micrometers per day in cultured rat hippocampal neurons [14]. Accordingly, we consider a range of values of *v*_e_ = 1 µm h^−1^ − 10 µm h^−1^, as reasonable such that the growth rate for a neurite accumulating most growth factors will be within 1 µm h^−1^ - 10 µm h^−1^.

#### Polarity and growth factor equilibrium constants

Reasonable values for the polarity and growth factors’ equilibrium constants, *K*_p_ and *K*_g_ respectively, should be within 1 to 100, based on the rest of the parametrization. These are the values compared to the number of bound MT for which Eq (32) and Eq (33) remain reasonable.

### Bifurcation analysis of the neuron model

We determine the parameter regimes under which our neuron model admits multiple axon-like neurites. Throughout this section, we first consider a neuron in which all neurites have equal and fixed length *L*, implemented by setting the growth speed *v*_e_ = 0. To simplify the analysis, we examine an ultrasensitive limit of the polarization mechanism in which the stabilization of boundary MTs becomes effectively switch-like. Mathematically, we take the Hill coefficient in the definition of *k*_u_ (Eq (34)) to infinity, *h*_u_ → ∞, yielding the Heaviside form

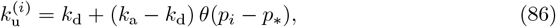

in which *θ* is the Heaviside step function and *i* indexes the neurites. In this limit, *k*_u_ behaves as a binary polarity selector: since 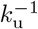 is the mean residence time in the boundary state, the two possible rates *k*_a_ and *k*_d_ correspond to two distinct steady-state numbers of boundary MTs, which we denote by *A* (axon-like) and *D* (dendrite-like), respectively. Thus, each neurite satisfies

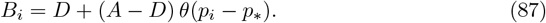

With *p*_*i*_ from Eq (32) we find the self-consistency condition

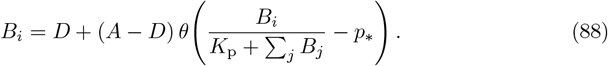

Suppose the steady state contains *n*_a_ axon-like neurites (*B* = *A*) and *n* − *n*_a_ dendrite-like neurites (*B* = *D*). Summing Eq (88) over all neurites yields a single self-consistency equation for the integer variable *n*_a_:

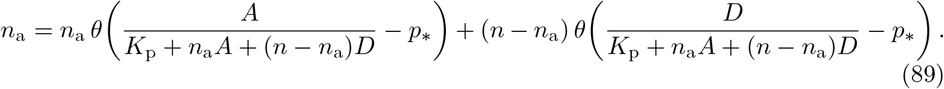

Rearranging this expression gives

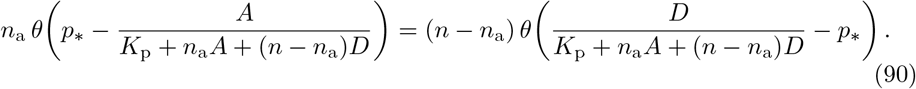

For a given parameter set, the integers *n*_a_ satisfying Eq (90) are the possible steady-state numbers of axon-like neurites. The all-dendrite state *n*_a_ = 0 exists when

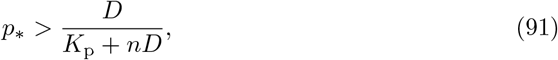

and the all-axon state *n*_a_ = *n* exists when

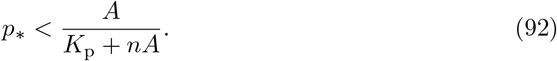

For intermediate values 1 ≤ *n*_a_ ≤ *n* − 1, the condition reduces to

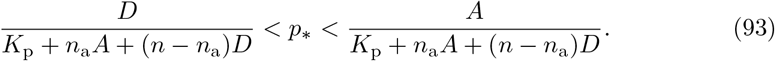

Because all neurites are identical in this scenario, any permutation of axonal identities yields an equivalent steady state, so each admissible value of *n*_a_ corresponds to a degenerate manifold of solutions.

To break this degeneracy, we now let one neurite be longer than the remaining *n* − 1, which remain equal in length. Length affects the number of boundary MTs through two competing mechanisms: longer neurites admit more non-centrosomal MTs nucleated in the neurite to reach the tip, whereas shorter neurites admit more centrosomal MTs from the cell soma. We consider the regime in which the first effect dominates, implying

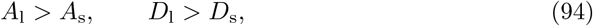

where subscripts l and s denote long and short neurites. A configuration is now specified by two integers: the number of axon-like short neurites *n*_a,*s*_ ∈ *{*0, …, *n* − 1} and whether the long neurite is axon-like, *n*_a,*l*_ ∈ {0, 1}. Summing the analogue of Eq (88) over all neurites yields

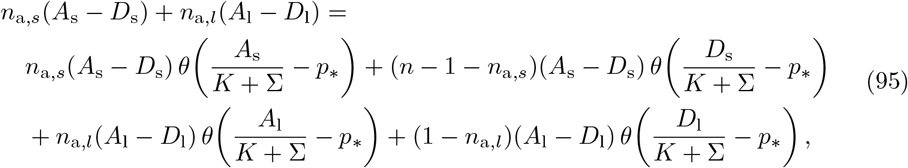

with

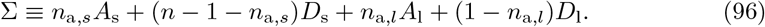

Introducing the ratio

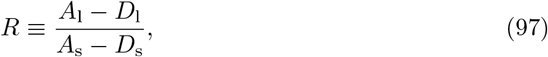

the condition simplifies to

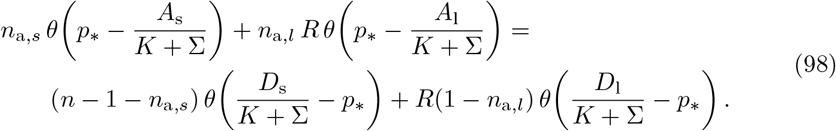

We focus on the biologically relevant regime in which the long neurite becomes the unique axon, i.e., *n*_a,*l*_ = 1 and *n*_a,*s*_ = 0, while all short neurites remain dendritic. The consistency condition then reduces to

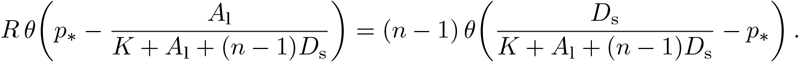

This equality is satisfied precisely when

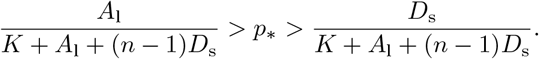

A second algebraic solution exists when *D*_s_ *> A*_l_ and *R* = *n ™* 1, but this contradicts the biological ordering *A*_l_ *> A*_s_ *> D*_s_ and is therefore discarded. To ensure that this solution is the only stable configuration, the inequalities required for competing states, such as *n*_a,*l*_ = 1 with *n*_a,*s*_ ≠ 0, or *n*_a,*l*_ = 0, must fail to hold. These statements can be verified directly using the general self-consistency equation above.

## Acknowledgments

K.N. acknowledges funding from the Trappeniers-Wols Fonds.

## Notes

### Competing Interest Statement

The authors have declared no competing interest.

## References

1. Sánchez-Huertas C, Herrera E. With the Permission of Microtubules: An Updated Overview on Microtubule Function During Axon Pathfinding. Frontiers in Molecular Neuroscience. 2021 Dec;14:759404. Available from: https://www.frontiersin.org/articles/10.3389/fnmol.2021.759404/full. doi:10.3389/fnmol.2021.759404.

2. Stiess M, Bradke F. Neuronal polarization: The cytoskeleton leads the way. Developmental Neurobiology. 2011 Jun;71(6):430–44. Available from: https://onlinelibrary.wiley.com/doi/10.1002/dneu.20849. doi:10.1002/dneu.20849.

3. Higgs VE, Das RM. Establishing neuronal polarity: microtubule regulation during neurite initiation. Oxford Open Neuroscience. 2022 May;1:kvac007. Available from: https://www.ncbi.nlm.nih.gov/pmc/articles/PMC10913830/. doi:10.1093/oons/kvac007.

4. Cyske Z, Gaffke L, Pierzynowska K, Węgrzyn G. Tubulin Cytoskeleton in Neurodegenerative Diseases–not Only Primary Tubulinopathies. Cellular and Molecular Neurobiology. 2023 Jul;43(5):1867–84. Available from: https://link.springer.com/10.1007/s10571-022-01304-6. doi:10.1007/s10571-022-01304-6.

5. Baas PW, Deitch JS, Black MM, Banker GA. Polarity orientation of microtubules in hippocampal neurons: uniformity in the axon and nonuniformity in the dendrite. Proceedings of the National Academy of Sciences. 1988 Nov;85(21):8335-9. Publisher: Proceedings of the National Academy of Sciences. Available from: https://www.pnas.org/doi/abs/10.1073/pnas.85.21.8335. doi: 10.1073/pnas.85.21.8335.

6. Yau KW, Schätzle P, Tortosa E, Pagès S, Holtmaat A, Kapitein LC, et al. Dendrites In Vitro and In Vivo Contain Microtubules of Opposite Polarity and Axon Formation Correlates with Uniform Plus-End-Out Microtubule Orientation. The Journal of Neuroscience. 2016 Jan;36(4):1071–85. Available from: https://www.jneurosci.org/lookup/doi/10.1523/JNEUROSCI.2430-15.2016. doi:10.1523/JNEUROSCI.2430-15.2016.

7. Binder LI, Frankfurter A, Rebhun LI. The distribution of tau in the mammalian central nervous system. Journal of Cell Biology. 1985 Oct;101(4):1371–8. Available from: https://doi.org/10.1083/jcb.101.4.1371. doi:10.1083/jcb.101.4.1371.

8. Kanai Y, Hirokawa N. Sorting mechanisms of Tau and MAP2 in neurons: Suppressed axonal transit of MAP2 and locally regulated microtubule binding. Neuron. 1995 Feb;14(2):421–32. Available from: https://linkinghub.elsevier.com/retrieve/pii/0896627395902988. doi:10.1016/0896-6273(95)90298-8.

9. Tortosa E, Adolfs Y, Fukata M, Pasterkamp RJ, Kapitein LC, Hoogenraad CC. Dynamic Palmitoylation Targets MAP6 to the Axon to Promote Microtubule Stabilization during Neuronal Polarization. Neuron. 2017 May;94(4):809-25.e7. Available from: https://linkinghub.elsevier.com/retrieve/pii/S0896627317303999. doi:10.1016/j.neuron.2017.04.042.

10. Monroy BY, Tan TC, Oclaman JM, Han JS, Simó S, Niwa S, et al. A Combinatorial MAP Code Dictates Polarized Microtubule Transport. Developmental Cell. 2020 Apr;53(1):60-72.e4. Publisher: Elsevier. Available from: https://www.cell.com/developmental-cell/abstract/S1534-5807(20)30061-7. doi:10.1016/j.devcel.2020.01.029.

11. van Beuningen SFB, Will L, Harterink M, Chazeau A, van Battum EY, Frias CP, et al. TRIM46 Controls Neuronal Polarity and Axon Specification by Driving the Formation of Parallel Microtubule Arrays. Neuron. 2015 Dec;88(6):1208–26. doi:10.1016/j.neuron.2015.11.012.

12. Tas RP, Chazeau A, Cloin BMC, Lambers MLA, Hoogenraad CC, Kapitein LC. Differentiation between Oppositely Oriented Microtubules Controls Polarized Neuronal Transport. Neuron. 2017 Dec;96(6):1264-71.e5. Available from: https://linkinghub.elsevier.com/retrieve/pii/S0896627317310711. doi:10.1016/j.neuron.2017.11.018.

13. Harterink M, Edwards SL, De Haan B, Yau KW, Van Den Heuvel S, Kapitein LC, et al. Local microtubule organization promotes cargo transport in C. elegans dendrites. Journal of Cell Science. 2018 Oct;131(20):jcs223107. Available from: https://journals.biologists.com/jcs/article/131/20/jcs223107/80/Local-microtubule-organization-promotes-cargo. doi:10.1242/jcs.223107.

14. Dotti C, Sullivan C, Banker G. The establishment of polarity by hippocampal neurons in culture. The Journal of Neuroscience. 1988 Apr;8(4):1454–68. Available from: https://www.jneurosci.org/lookup/doi/10.1523/JNEUROSCI.08-04-01454.1988. doi:10.1523/JNEUROSCI.08-04-01454.1988.

15. Ahmad FJ, Echeverri CJ, Vallee RB, Baas PW. Cytoplasmic Dynein and Dynactin Are Required for the Transport of Microtubules into the Axon. Journal of Cell Biology. 1998 Jan;140(2):391–401. Available from: https://doi.org/10.1083/jcb.140.2.391. doi:10.1083/jcb.140.2.391.

16. Burute M, Jansen KI, Mihajlovic M, Vermonden T, Kapitein LC. Local changes in microtubule network mobility instruct neuronal polarization and axon specification. Science Advances. 2022 Nov;8(44):eabo2343. Publisher: American Association for the Advancement of Science. Available from: https://www.science.org/doi/full/10.1126/sciadv.abo2343. doi:10.1126/sciadv.abo2343.

17. Oelz DB, Del Castillo U, Gelfand VI, Mogilner A. Microtubule Dynamics, Kinesin-1 Sliding, and Dynein Action Drive Growth of Cell Processes. Biophysical Journal. 2018 Oct;115(8):1614–24. Available from: https://linkinghub.elsevier.com/retrieve/pii/S0006349518310245. doi:10.1016/j.bpj.2018.08.046.

18. Guha S, Patil A, Muralidharan H, Baas PW. Mini-review: Microtubule sliding in neurons. Neuroscience Letters. 2021 May;753:135867. Available from: https://www.sciencedirect.com/science/article/pii/S0304394021002457. doi:10.1016/j.neulet.2021.135867.

19. Sánchez-Huertas C, Freixo F, Viais R, Lacasa C, Soriano E, Lüders J. Non-centrosomal nucleation mediated by augmin organizes microtubules in post-mitotic neurons and controls axonal microtubule polarity. Nature Communications. 2016 Jul;7(1):12187. Publisher: Nature Publishing Group. Available from: https://www.nature.com/articles/ncomms12187. doi:10.1038/ncomms12187.

20. Cunha-Ferreira I, Chazeau A, Buijs RR, Stucchi R, Will L, Pan X, et al. The HAUS Complex Is a Key Regulator of Non-centrosomal Microtubule Organization during Neuronal Development. Cell Reports. 2018 Jul;24(4):791–800. Available from: https://www.sciencedirect.com/science/article/pii/S2211124718310350. doi:10.1016/j.celrep.2018.06.093.

21. Zhang Y, Sung HH, Ziegler AB, Wu YC, Viais R, Sánchez-Huertas C, et al. Augmin complex activity finetunes dendrite morphology through non-centrosomal microtubule nucleation in vivo. Journal of Cell Science. 2024 May;137(9):jcs261512. Available from: https://www.ncbi.nlm.nih.gov/pmc/articles/PMC11128282/. doi:10.1242/jcs.261512.

22. Tymanskyj SR, Yang BH, Verhey KJ, Ma L. MAP7 regulates axon morphogenesis by recruiting kinesin-1 to microtubules and modulating organelle transport. eLife. 2018 Aug;7:e36374. Publisher: eLife Sciences Publications, Ltd. Available from: https://doi.org/10.7554/eLife.36374. doi:10.7554/eLife.36374.

23. Harterink M, Vocking K, Pan X, Soriano Jerez EM, Slenders L, Fréal A, et al. TRIM46 Organizes Microtubule Fasciculation in the Axon Initial Segment. The Journal of Neuroscience. 2019 Jun;39(25):4864–73. Available from: https://www.ncbi.nlm.nih.gov/pmc/articles/PMC6670255/. doi:10.1523/JNEUROSCI.3105-18.2019.

24. Witte H, Neukirchen D, Bradke F. Microtubule stabilization specifies initial neuronal polarization. The Journal of Cell Biology. 2008 Feb;180(3):619–32. Available from: https://rupress.org/jcb/article/180/3/619/34944/Microtubule-stabilization-specifies-initial. doi:10.1083/jcb.200707042.

25. Gomis-Rüth S, Wierenga CJ, Bradke F. Plasticity of Polarization: Changing Dendrites into Axons in Neurons Integrated in Neuronal Circuits. Current Biology. 2008 Jul;18(13):992-1000. Publisher: Elsevier. Available from: https://www.cell.com/current-biology/abstract/S0960-9822(08)00748-3. doi:10.1016/j.cub.2008.06.026.

26. Scanlon HG, Mahata G, Nelson AC, McKinley SA, Rolls MM, Ciocanel MV. Nucleation feedback can drive establishment and maintenance of biased microtubule polarity in neurites. Mathematical Biosciences. 2025 Nov;389:109538. Available from: https://linkinghub.elsevier.com/retrieve/pii/S0025556425001646. doi:10.1016/j.mbs.2025.109538.

27. Samuels DC, Hentschel HGE, Fine A. The origin of neuronal polarization: a model of axon formation. Philosophical Transactions of the Royal Society of London Series B: Biological Sciences. 1996 Sep;351(1344):1147–56. Available from: https://royalsocietypublishing.org/rstb/article/351/1344/1147/18934/The-origin-of-neuronal-polarization-a-model-of. doi:10.1098/rstb.1996.0099.

28. Van Ooyen A. Using theoretical models to analyse neural development. Nature Reviews Neuroscience. 2011 Jun;12(6):311–26. Available from: https://www.nature.com/articles/nrn3031. doi:10.1038/nrn3031.

29. Shelly M, Cancedda L, Lim BK, Popescu A, Cheng Pl, Gao H, et al. Semaphorin3A Regulates Neuronal Polarization by Suppressing Axon Formation and Promoting Dendrite Growth. Neuron. 2011 Aug;71(3):433–46. Available from: https://www.ncbi.nlm.nih.gov/pmc/articles/PMC3164872/. doi:10.1016/j.neuron.2011.06.041.

30. Förster E. Reelin, neuronal polarity and process orientation of cortical neurons. Neuroscience. 2014 Jun;269:102–11. Available from: https://www.sciencedirect.com/science/article/pii/S0306452214002103. doi:10.1016/j.neuroscience.2014.03.004.

31. Puri D, Ponniah K, Biswas K, Basu A, Dey S, Lundquist EA, et al. Wnt signaling establishes the microtubule polarity in neurons through regulation of Kinesin-13. Journal of Cell Biology. 2021 Sep;220(9):e202005080. Available from: https://rupress.org/jcb/article/220/9/e202005080/212396/Wnt-signaling-establishes-the-microtubule-polarity. doi:10.1083/jcb.202005080.

32. Koser DE, Thompson AJ, Foster SK, Dwivedy A, Pillai EK, Sheridan GK, et al. Mechanosensing is critical for axon growth in the developing brain. Nature neuroscience. 2016 Dec;19(12):1592–8. Available from: https://www.ncbi.nlm.nih.gov/pmc/articles/PMC5531257/. doi:10.1038/nn.4394.

33. Schelski M, Bradke F. Neuronal polarization: From spatiotemporal signaling to cytoskeletal dynamics. Molecular and Cellular Neuroscience. 2017 Oct;84:11–28. Available from: https://www.sciencedirect.com/science/article/pii/S1044743116302561. doi:10.1016/j.mcn.2017.03.008.

34. De Rooij R, Kuhl E, Miller KE. Modeling the Axon as an Active Partner with the Growth Cone in Axonal Elongation. Biophysical Journal. 2018 Nov;115(9):1783–95. Available from: https://linkinghub.elsevier.com/retrieve/pii/S0006349518310610. doi:10.1016/j.bpj.2018.08.047.

35. Oliveri H, Goriely A. Mathematical models of neuronal growth. Biomechanics and Modeling in Mechanobiology. 2022;21(1):89–118. Available from: https://www.ncbi.nlm.nih.gov/pmc/articles/PMC8807500/. doi:10.1007/s10237-021-01539-0.

36. Jakobs MA, Zemel A, Franze K. Unrestrained growth of correctly oriented microtubules instructs axonal microtubule orientation. eLife. 2022 Oct;11:e77608. Available from: https://elifesciences.org/articles/77608. doi:10.7554/eLife.77608.

37. Takano T, Wu M, Nakamuta S, Naoki H, Ishizawa N, Namba T, et al. Discovery of long-range inhibitory signaling to ensure single axon formation. Nature Communications. 2017 Jun;8(1):33. Publisher: Nature Publishing Group. Available from: https://www.nature.com/articles/s41467-017-00044-2. doi:10.1038/s41467-017-00044-2.

38. Toriyama M, Sakumura Y, Shimada T, Ishii S, Inagaki N. A diffusion-based neurite length-sensing mechanism involved in neuronal symmetry breaking. Molecular Systems Biology. 2010 Jan;6(1):394. Available from: https://link.springer.com/article/10.1038/msb.2010.51. doi:10.1038/msb.2010.51.

39. Hall A, Lalli G. Rho and Ras GTPases in Axon Growth, Guidance, and Branching. Cold Spring Harbor Perspectives in Biology. 2010 Feb;2(2):a001818–8. Available from: http://cshperspectives.cshlp.org/lookup/doi/10.1101/cshperspect.a001818. doi:10.1101/cshperspect.a001818.

40. Fivaz M, Bandara S, Inoue T, Meyer T. Robust Neuronal Symmetry Breaking by Ras-Triggered Local Positive Feedback. Current Biology. 2008 Jan;18(1):44–50. Available from: https://linkinghub.elsevier.com/retrieve/pii/S0960982207023366. doi:10.1016/j.cub.2007.11.051.

41. Govek EE, Newey SE, Aelst LV. The role of the Rho GTPases in neuronal development. Genes & Development. 2005 Jan;19(1):1–49. Company: Cold Spring Harbor Laboratory Press Distributor: Cold Spring Harbor Laboratory Press Institution: Cold Spring Harbor Laboratory Press Label: Cold Spring Harbor Laboratory Press Publisher: Cold Spring Harbor Lab. Available from: http://genesdev.cshlp.org/content/19/1/1. doi:10.1101/gad.1256405.

42. Foteinopoulos P, Mulder BM. A microtubule-based minimal model for spontaneous and persistent spherical cell polarity. PLOS ONE. 2017 Sep;12(9):e0184706. Publisher: Public Library of Science. Available from: https://journals.plos.org/plosone/article?id=10.1371/journal.pone.0184706. doi:10.1371/journal.pone.0184706.

43. Meinhardt H. Orientation of chemotactic cells and growth cones: models and mechanisms. Journal of Cell Science. 1999 Sep;112(17):2867–74. Available from: https://doi.org/10.1242/jcs.112.17.2867. doi:10.1242/jcs.112.17.2867.

44. Rao AN, Patil A, Black MM, Craig EM, Myers KA, Yeung HT, et al. Cytoplasmic Dynein Transports Axonal Microtubules in a Polarity-Sorting Manner. Cell Reports. 2017 Jun;19(11):2210–9. Available from: https://linkinghub.elsevier.com/retrieve/pii/S2211124717307210. doi:10.1016/j.celrep.2017.05.064.

45. Winding M, Kelliher MT, Lu W, Wildonger J, Gelfand VI. Role of kinesin-1–based microtubule sliding in Drosophila nervous system development. Proceedings of the National Academy of Sciences. 2016 Aug;113(34). Available from: https://pnas.org/doi/full/10.1073/pnas.1522416113. doi:10.1073/pnas.1522416113.

46. Yan J, Chao DL, Toba S, Koyasako K, Yasunaga T, Hirotsune S, et al. Kinesin-1 regulates dendrite microtubule polarity in Caenorhabditis elegans. eLife. 2013 Mar;2:e00133. Publisher: eLife Sciences Publications, Ltd. Available from: https://doi.org/10.7554/eLife.00133. doi:10.7554/eLife.00133.

47. Sharp DJ, Ross JL. Microtubule-severing enzymes at the cutting edge. Journal of Cell Science. 2012 Jun;125(11):2561–9. Available from: https://www.ncbi.nlm.nih.gov/pmc/articles/PMC3403230/. doi:10.1242/jcs.101139.

48. Akhmanova A, Kapitein LC. Mechanisms of microtubule organization in differentiated animal cells. Nature Reviews Molecular Cell Biology. 2022 Aug;23(8):541–58. Available from: https://www.nature.com/articles/s41580-022-00473-y. doi:10.1038/s41580-022-00473-y.

49. Rao AN, Baas PW. Polarity Sorting of Microtubules in the Axon. Trends in Neurosciences. 2018 Feb;41(2):77–88. Publisher: Elsevier. Available from: https://www.cell.com/trends/neurosciences/abstract/S0166-2236(17)30214-X. doi:10.1016/j.tins.2017.11.002.

50. Ahmad FJ, Pienkowski TP, Baas PW. Regional differences in microtubule dynamics in the axon. Journal of Neuroscience. 1993 Feb;13(2):856–66. Publisher: Society for Neuroscience Section: Articles. Available from: https://www.jneurosci.org/content/13/2/856. doi:10.1523/JNEUROSCI.13-02-00856.1993.

51. Prezel E, Elie A, Delaroche J, Stoppin-Mellet V, Bosc C, Serre L, et al. Tau can switch microtubule network organizations: from random networks to dynamic and stable bundles. Molecular Biology of the Cell. 2018 Jan;29(2):154–65. Publisher: American Society for Cell Biology (mboc). Available from: https://www.molbiolcell.org/doi/full/10.1091/mbc.E17-06-0429. doi:10.1091/mbc.E17-06-0429.

52. Costa AC, Sousa MM. The Role of Spastin in Axon Biology. Frontiers in Cell and Developmental Biology. 2022 Jul;10. Publisher: Frontiers. Available from: https://www.frontiersin.org/journals/cell-and-developmental-biology/articles/10.3389/fcell.2022.934522/full. doi:10.3389/fcell.2022.934522.

53. Sarbanes SL, Zehr EA, Roll-Mecak A. Microtubule-severing enzymes. Current Biology. 2022 Oct;32(19):R992–7. Publisher: Elsevier. Available from: https://www.cell.com/current-biology/abstract/S0960-9822(22)01368-9. doi:10.1016/j.cub.2022.08.046.

54. Wilkes OR, Moore AW. Distinct Microtubule Organizing Center Mechanisms Combine to Generate Neuron Polarity and Arbor Complexity. Frontiers in Cellular Neuroscience. 2020 Nov;14:594199. Available from: https://www.frontiersin.org/articles/10.3389/fncel.2020.594199/full. doi:10.3389/fncel.2020.594199.

55. Petry S, Groen A, Ishihara K, Mitchison T, Vale R. Branching Microtubule Nucleation in Xenopus Egg Extracts Mediated by Augmin and TPX2. Cell. 2013 Feb;152(4):768–77. Available from: https://linkinghub.elsevier.com/retrieve/pii/S0092867413000159. doi:10.1016/j.cell.2012.12.044.

56. Twelvetrees AE, Pernigo S, Sanger A, Guedes-Dias P, Schiavo G, Steiner RA, et al. The Dynamic Localization of Cytoplasmic Dynein in Neurons Is Driven by Kinesin-1. Neuron. 2016 Jun;90(5):1000–15. Publisher: Elsevier. Available from: https://www.cell.com/neuron/abstract/S0896-6273(16)30160-X. doi:10.1016/j.neuron.2016.04.046.

57. Dogterom M, Leibler S. Physical aspects of the growth and regulation of microtubule structures. Physical Review Letters. 1993 Mar;70(9):1347–50. Publisher: American Physical Society. Available from: https://link.aps.org/doi/10.1103/PhysRevLett.70.1347. doi:10.1103/PhysRevLett.70.1347.

58. Jacobson C, Schnapp B, Banker GA. A Change in the Selective Translocation of the Kinesin-1 Motor Domain Marks the Initial Specification of the Axon. Neuron. 2006 Mar;49(6):797–804. Available from: https://linkinghub.elsevier.com/retrieve/pii/S089662730600095X. doi:10.1016/j.neuron.2006.02.005.

59. Toriyama M, Shimada T, Kim KB, Mitsuba M, Nomura E, Katsuta K, et al. Shootin1: a protein involved in the organization of an asymmetric signal for neuronal polarization. The Journal of Cell Biology. 2006 Oct;175(1):147–57. Available from: https://rupress.org/jcb/article/175/1/147/34344/Shootin1-a-protein-involved-in-the-organization-of. doi:10.1083/jcb.200604160.

60. Schelski M, Bradke F. Microtubule retrograde flow retains neuronal polarization in a fluctuating state. Science Advances. 2022 Nov;8(44):eabo2336. Available from: https://www.science.org/doi/10.1126/sciadv.abo2336. doi:10.1126/sciadv.abo2336.

61. Iwanski MK, Serweta AK, Van Schelt J, Verwei HN, Donders BC, Kapitein LC. Polarity reversal of stable microtubules during neuronal development. Journal of Cell Science. 2025 Nov;138(22):jcs264152. Available from: https://journals.biologists.com/jcs/article/138/22/jcs264152/369969/Polarity-reversal-of-stable-microtubules-during. doi:10.1242/jcs.264152.

62. Baas PW, Black MM, Banker GA. Changes in microtubule polarity orientation during the development of hippocampal neurons in culture. The Journal of cell biology. 1989 Dec;109(6):3085–94. Available from: https://rupress.org/jcb/article/109/6/3085/29006/Changes-in-microtubule-polarity-orientation-during. doi:10.1083/jcb.109.6.3085.

63. Kuo YW, Howard J. Cutting, Amplifying, and Aligning Microtubules with Severing Enzymes. Trends in Cell Biology. 2021 Jan;31(1):50–61. Available from: https://www.cell.com/trends/cell-biology/abstract/S0962-8924(20)30210-5. doi:10.1016/j.tcb.2020.10.004.

64. Lowery LA, Vactor DV. The trip of the tip: understanding the growth cone machinery. Nature Reviews Molecular Cell Biology. 2009 May;10(5):332–43. Available from: https://www.nature.com/articles/nrm2679. doi:10.1038/nrm2679.

65. Coles C, Bradke F. Coordinating Neuronal Actin–Microtubule Dynamics. Current Biology. 2015 Aug;25(15):R677–91. Available from: https://linkinghub.elsevier.com/retrieve/pii/S0960982215007149. doi:10.1016/j.cub.2015.06.020.

66. Baas PW, Rao AN, Matamoros AJ, Leo L. Stability properties of neuronal microtubules. Cytoskeleton. 2016 Sep;73(9):442–60. Available from: https://onlinelibrary.wiley.com/doi/10.1002/cm.21286. doi:10.1002/cm.21286.

67. Iwanski MK, Kapitein LC. Cellular cartography: Towards an atlas of the neuronal microtubule cytoskeleton. Frontiers in Cell and Developmental Biology. 2023 Mar;11:1052245. Available from: https://www.frontiersin.org/articles/10.3389/fcell.2023.1052245/full. doi:10.3389/fcell.2023.1052245.

68. Gillespie DT. Stochastic Simulation of Chemical Kinetics. Annual Review of Physical Chemistry. 2007 May;58(1):35–55. Available from: https://www.annualreviews.org/doi/10.1146/annurev.physchem.58.032806.104637. doi:10.1146/annurev.physchem.58.032806.104637.

69. Yogev S, Cooper R, Fetter R, Horowitz M, Shen K. Microtubule organization determines axonal transport dynamics. Neuron. 2016 Oct;92(2):449–60. Available from: https://www.ncbi.nlm.nih.gov/pmc/articles/PMC5432135/. doi:10.1016/j.neuron.2016.09.036.

70. Yu W, Baas PW. Changes in microtubule number and length during axon differentiation. Journal of Neuroscience. 1994 May;14(5):2818–29. Publisher: Society for Neuroscience Section: Articles. Available from: https://www.jneurosci.org/content/14/5/2818. doi:10.1523/JNEUROSCI.14-05-02818.1994.

71. Wang J, Yu W, Baas PW, Black MM. Microtubule Assembly in Growing Dendrites. Journal of Neuroscience. 1996 Oct;16(19):6065–78. Publisher: Society for Neuroscience Section: Articles. Available from: https://www.jneurosci.org/content/16/19/6065. doi:10.1523/JNEUROSCI.16-19-06065.1996.

72. Kapitein L, Hoogenraad C. Building the Neuronal Microtubule Cytoskeleton. Neuron. 2015 Aug;87(3):492–506. Available from: https://linkinghub.elsevier.com/retrieve/pii/S0896627315005139. doi:10.1016/j.neuron.2015.05.046.

73. Katrukha EA, Jurriens D, Salas Pastene DM, Kapitein LC. Quantitative mapping of dense microtubule arrays in mammalian neurons. eLife. 2021 Jul;10:e67925. Publisher: eLife Sciences Publications, Ltd. Available from: https://doi.org/10.7554/eLife.67925. doi:10.7554/eLife.67925.

74. Wang L, Brown A. Rapid Movement of Microtubules in Axons. Current Biology. 2002 Sep;12(17):1496–501. Available from: https://www.sciencedirect.com/science/article/pii/S0960982202010783. doi:10.1016/S0960-9822(02)01078-3.

